# Engineering the NAT–ULT module increases seed number per pod in soybean

**DOI:** 10.64898/2026.10.09.758010

**Authors:** Cuong X. Nguyen, Jae hyo Song, Vikranth Kumar, Manh V. Nguyen, Sterling Evans, Shunhong Dai, Minviluz G. Stacey

## Abstract

Seed number per pod is a major soybean yield-component trait, yet experimentally validated causal modules remain poorly defined. Here, we identify and functionally validate a soybean NAT–ULT regulatory module controlling ovule and seed number per pod. Through a visual screen of a fast-neutron-irradiated Glycine max cv. Williams 82 population, we isolated MO27, which increased four-seeded pod frequency from approximately 2–4% in wild type to 27–30%, without major changes in plant architecture, total pod number, or seed morphology. Comparative genome hybridization mapped the phenotype to a 46.7-kb chromosome 2 deletion containing *GmNAT1* and chromosome 6 deletions encompassing *GmULT-B1* and *GmULT-B3*. Ovule-clearing analysis showed that increased seed number originated before fertilization through elevated formation of four-ovule ovaries. CRISPR/Cas9-mediated mutagenesis of *GmNAT1*, *GmULT-B1*, and *GmULT-B3* increased four-seeded pod frequency without detectable penalties in 100-seed weight, total pod number, or plant height. Tomato *SlULT* triple mutants also increased floral organ and locule number, supporting conserved ULT-like function in reproductive organ patterning. RNA-seq interaction analysis revealed a chromosome 6-dominant transcriptional state, accompanied by selective non-additive reprogramming of hormone/ovule-number and cell-wall-remodeling modules. Integrated genetic and transcriptomic analyses support a hierarchical model in which NAT–ULT perturbation may enhance meristematic competence and remodel reproductive developmental programs, while additional fast-neutron-background-associated transcriptional changes may further modulate the phenotype. These findings define NAT–ULT as a molecular target for engineering soybean seed number per pod.

## Introduction

Soybean (*Glycine max* [L.] Merr.) is one of the most important legume crops worldwide, providing a major source of plant protein and vegetable oil for food, feed, and industrial uses. Continued growth in global demand has intensified the need to improve soybean yield potential and yield stability. Soybean yield is determined by the combined effects of plant density, pod number per plant, seed number per pod, and individual seed weight, with seed number per unit area generally representing the most important yield component (Ainsworth *et al*., 2012; Vogel *et al*., 2021). Although breeding and genetic studies have substantially improved traits related to plant architecture, flowering time, pod production, and seed filling, seed number per pod remains a comparatively underexploited yield component. This limitation partly reflects the rarity of naturally occurring soybean variants with consistently increased seed number per pod and the limited mechanistic understanding of the genetic pathways that control this trait (Pedersen and Lauer, 2004)

In soybean, most pods contain one to three seeds, whereas four-seeded pods usually represent only a small fraction of total pod output in cultivated varieties (Yang *et al*., 2013). Because the maximum seed number per pod is constrained by the number of ovules initiated in the ovary before fertilization, increasing ovule number provides a direct developmental route to increasing seed number. Ovule number is established during early reproductive development through coordinated regulation of placenta growth, auxin distribution, boundary formation, and ovule primordium initiation (Cucinotta *et al*., 2014, 2020; Galbiati *et al*., 2013; Klucher *et al*., 1996; Mizukami and Fischer, 2000). Therefore, genes that regulate ovule initiation, floral meristem activity, or placenta-derived organ formation represent attractive targets for improving seed number per pod. However, despite recent fine-mapping, GWAS, and QTN studies implicating multiple genomic regions, candidate genes, and marker-assisted selection models for soybean seed number per pod and four-seeded pod traits, the causal genes and developmental mechanisms that directly regulate ovule number remain poorly defined (Li *et al*., 2021; Wang *et al*., 2024; Yang *et al*., 2013; Yuan *et al*., 2025).

Studies in model plants have shown that reproductive organ number is controlled by regulatory networks that balance meristem maintenance, organ founder-cell specification, auxin distribution, and floral meristem determinacy. The CLAVATA–WUSCHEL (CLV-WUS) pathway (Clark *et al*., 1996), *SHOOTMERISTEMLESS*-related meristem regulators (Scofield *et al*., 2007), *CUP-SHAPED COTYLEDON* boundary genes (Galbiati *et al*., 2013), *AINTEGUMENTA*-mediated cell proliferation (Klucher *et al*., 1996), and TCP transcription factors all contribute to the control of lateral organ initiation or ovule development in Arabidopsis and other species (Cubas *et al*., 1999; Koyama *et al*., 2007). These pathways demonstrate that modest changes in meristem size, boundary formation, or organ identity can alter the number of carpels, locules, ovules, or seeds. Nevertheless, the extent to which these conserved developmental modules are connected to induced variation in soybean seed number per pod is still unclear.

ULTRAPETALA genes provide one potential link between meristem regulation and reproductive organ number. In Arabidopsis, *ULT1* and *ULT2* encode plant-specific SAND-domain proteins that influence floral meristem determinacy, organ number, and developmental gene expression (Carles *et al*., 2005; Fletcher, 2001). Loss of *ULT1* function increases floral meristem size and floral organ number, in part through altered regulation of the WUS–AGAMOUS (AG) network (Carles *et al*., 2005; Carles and Fletcher, 2009). ULT1 has also been described as a trithorax-group-associated regulator that promotes developmental gene expression through chromatin-level regulation (Carles and Fletcher, 2009). These findings place ULT proteins within a chromatin-associated regulatory layer that modulates reproductive meristem activity. However, whether ULT-related genes function in crop reproductive architecture, and whether they contribute to ovule or seed number variation in legumes, has remained unresolved.

Forward genetic screens using fast neutron (FN) mutagenesis offer a complementary strategy for identifying regulators of complex developmental traits in soybean. FN irradiation frequently generates chromosomal deletions or structural rearrangements that can disrupt single genes or linked gene clusters (Bolon *et al*., 2011). This feature is particularly useful in soybean, whose paleopolyploid genome contains extensive paralog retention and functional redundancy that can mask single-gene loss-of-function phenotypes (Schmutz *et al*., 2010). Screening FN populations for altered pod and seed traits can therefore reveal genetic modules that are difficult to detect through natural variation or single-gene reverse genetics alone.

Here, we identify a soybean NAT–ULT regulatory module that controls ovule and seed number per pod. Through a visual screen of a fast neutron-mutagenized Williams 82 population, we isolated the MO27 mutant, which displays a strong increase in four-seeded pod frequency. Genetic and molecular analyses linked this phenotype to deletion intervals containing *GmNAT1* and ULT-related genes including *GmULT-B1* and *GmULT-B3*. Targeted CRISPR/Cas9 mutagenesis confirmed that disruption of these genes increases four-seeded pod production, supporting a functional role for the NAT–ULT module in soybean reproductive development. Additional ovule-clearing, cross-species tomato validation, and transcriptomic analyses suggest that this module acts upstream of conserved meristem, boundary, and reproductive organ-patterning pathways. Together, these findings define a genetic entry point for increasing seed number per pod in soybean and provide a framework for connecting conserved floral developmental regulators with crop yield-component improvement.

## Result

### Identification of a FN–Induced Soybean Mutant with Increased Ovule Number and 4-SP

As part of a previously described visual screen of a FN–irradiated Glycine max cv. Williams 82 population (Stacey et al., 2016), we identified MO27 as a mutant line with altered pod-type distribution. MO27 was distinguished by a substantial increase in the frequency of 4-SP relative to WT. Representative pod images revealed a clear shift in pod composition in MO27, with an increased proportion of 4-SP accompanied by a corresponding reduction in two- and three-seeded pods (Figure 1a). Quantitative analysis confirmed that 4-SP were rare in WT, accounting for approximately 2–4% of total pods, whereas their frequency increased to approximately 27–30% in MO27 (Figure 1b).

**Figure 1.**
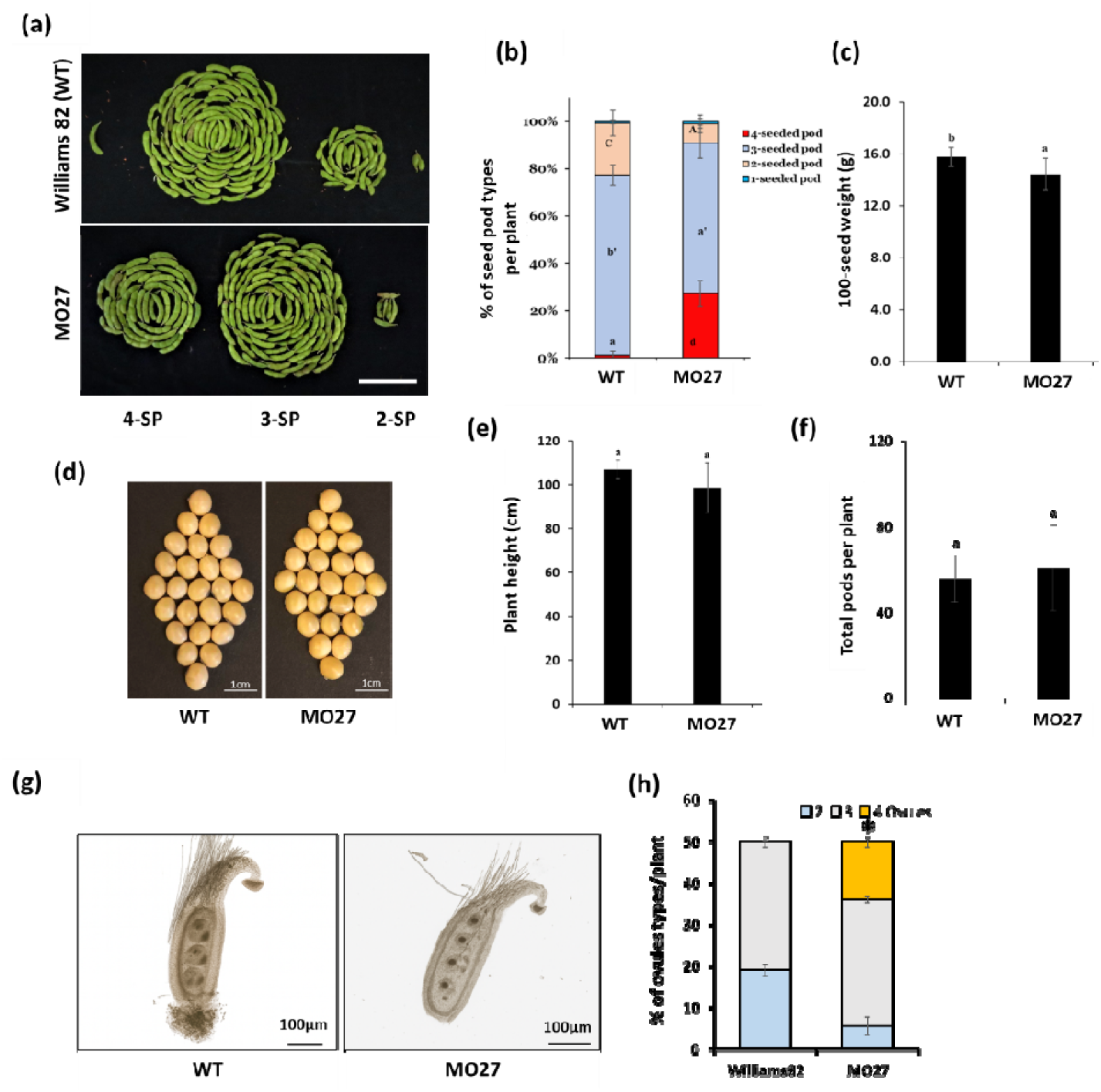
Identification and phenotypic characterization of a fast neutron-induced soybean mutant with an increased proportion of four-seeded pods. (a) Representative images showing pod type distribution in wild-type Glycine max cv. William 82 WT and the fast neutron-induced mutant line MO27, including four-seeded pods 4-SP, three-seeded pods 3-SP, and two-seeded pods 2-SP. Scale bar = 5 cm. (b) Proportion of pod types per plant in WT and MO27. MO27 exhibits a marked increase in the proportion of four-seeded pods relative to WT. Data are presented as mean ± SD, n = 20 plants. Different letters indicate significant differences as determined by one-way ANOVA followed by Tukey’s multiple comparison test P < 0.01. (c) Hundred-seed weight in WT and MO27. MO27 shows a modest reduction in seed weight relative to WT. Data are mean ± SD, n = 20 plants. (d) Representative seed morphology of WT and MO27. Scale bar = 1 cm. (e) Plant height and (f) total pod number per plant in WT and MO27. Data are mean ± SD n = 20 plants. (g) Representative cleared floral buds showing ovule number per ovary in WT and MO27. Scale bar = 100 µm. (h) Quantification of ovule number per ovary in WT and MO27. MO27 showed a significant shift in ovule-number distribution, including an increased proportion of four-ovule ovaries compared with WT. Data are presented as proportions from n = 100 ovaries per genotype. Statistical significance for the overall ovule-number distribution was determined using Pearson’s χ² test.

The increased frequency of 4-SP in MO27 occurred without broad changes in plant architecture or seed morphology. MO27 showed a modest but significant reduction in hundred-seed weight (Figure 1c), but representative seeds did not exhibit discernible differences in external morphology or overall shape compared with WT (Figure 1d). Likewise, plant height was not significantly altered (Figure 1e), and total pod number per plant remained comparable between genotypes (Figure 1f). Thus, the primary phenotypic effect of MO27 is an altered distribution of seed number per pod, rather than a generalized defect in vegetative growth, pod production, or seed morphology.

To determine whether the increased 4-SP phenotype originated before fertilization, we examined ovule number in pistils from unopened floral buds. Cleared ovary preparations showed that WT pistils predominantly contained two or three ovules, whereas MO27 frequently developed four ovules within a single ovary (Figure 1g). This difference was supported by quantitative analysis, which demonstrated a marked increase in the proportion of four-ovule ovaries in MO27 relative to WT (Figure 1h). In contrast, ovule length and width were not significantly different between genotypes (Figure S1), indicating that MO27 affects ovule number rather than individual ovule growth. The close correspondence between the frequency of four-ovule ovaries and mature 4-SP suggests that the MO27 phenotype is established during early floral development and is subsequently maintained through pod maturation.

### Comparative genome hybridization (CGH) and genetic analyses identify *GmNAT* and a chromosome 6 *GmULT1/2* locus associated with the four-seeded pod phenotype

To define the genetic basis of the elevated four-seeded pod frequency in MO27, we performed comparative genome hybridization (CGH) using wild-type Wm82 as the reference. CGH identified three FN-induced deletions in MO27, including a 46.7 kb deletion on chromosome 2 encompassing two predicted genes and two deletions on chromosome 6 spanning 224.5 kb and 651.0 kb, respectively (Figure 2a; Table S1).

**Figure 2.**
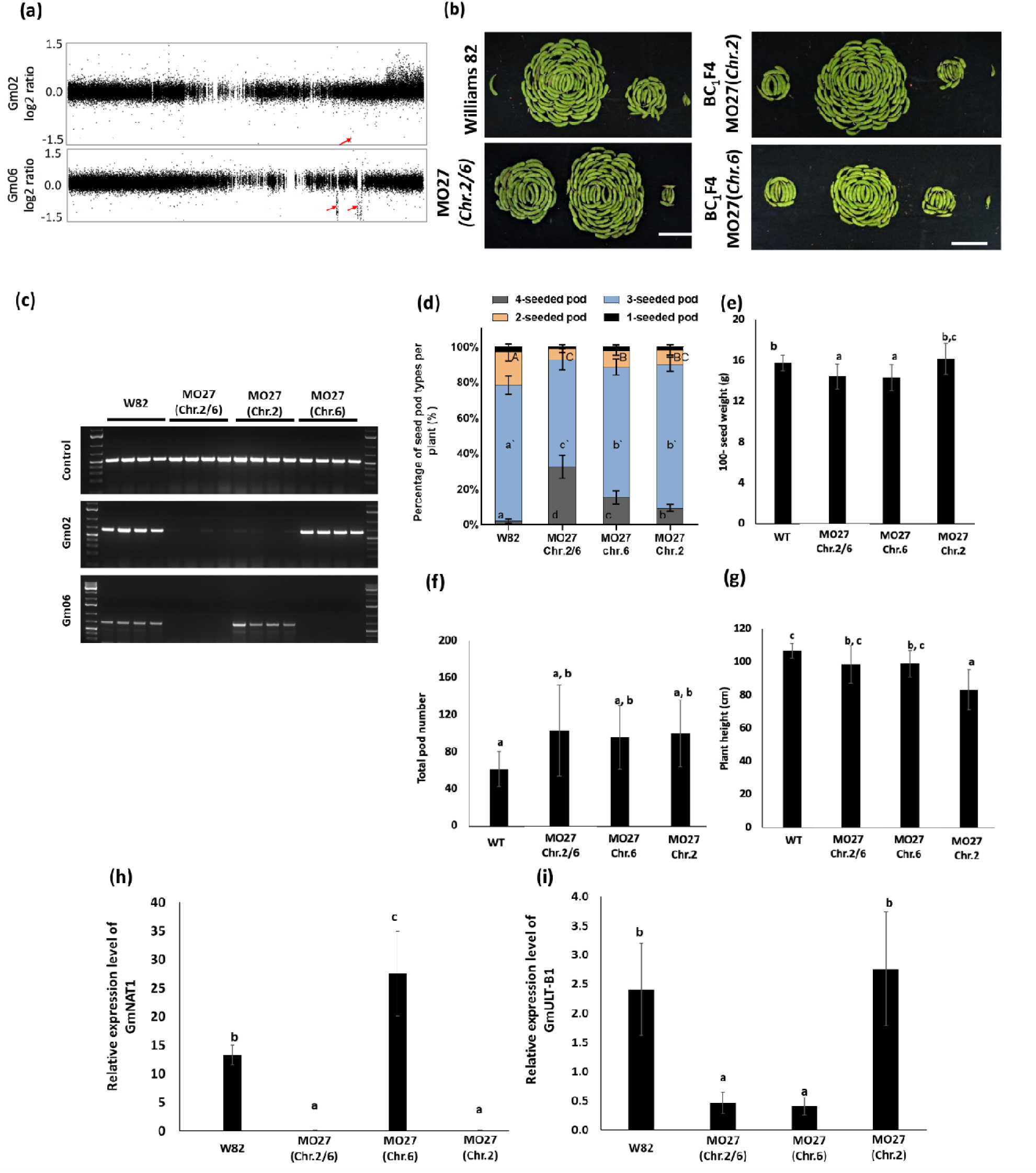
Genetic and phenotypic characterization of MO27-associated deletions underlying the increased four-seeded pod phenotype. (a) Comparative genome hybridization (CGH) profiles of MO27 relative to wild-type Glycine max cv. Williams 82 (WT), revealing deletions on chromosomes 2 and 6. Red arrows indicate deleted regions. (b) Representative pod phenotypes of WT, MO27, and BC_1_F4 plants segregating for chromosome 2 (chr.02) and chromosome 6 (chr.06) deletions. Scale bar = 5 cm. (c) PCR-based genotyping confirming the presence or absence of the chromosome 2 and chromosome 6 deletions in WT and segregating MO27-derived lines. Control indicates amplification using a Glyma.10G000700-specific primer set. Gm02 represents a GmNAT1-specific primer set for detection of the chromosome 2 deletion, and Gm06 represents a GmULT-B1-specific genotyping primer set for detection of the chromosome 6 deletion. (d) Proportion of pod types per plant in WT and deletion genotypes. Data are presented as mean ± SD (n = 20 plants). Different letters indicate statistically significant differences among genotypes (P < 0.01). (e–g) The 100-seed weight (e), total pod number per plant (f), and plant height (g) in WT and deletion genotypes. Data are presented as mean ± SD (n = 10 plants). (h, i) Relative expression levels of candidate genes located within the deleted regions in WT, MO27, and single-deletion lines (chr.2 and chr.6), determined by qRT-PCR. Data are presented as mean ± SD (n = 5). Statistical significance was assessed by one-way ANOVA followed by Tukey’s multiple range test, with different letters indicating significant differences (P < 0.01).

To determine which lesions contributed to the phenotype, MO27 was backcrossed, and 4-seeded pod frequency was assessed in the resulting BC_1_F2 populations. In the BC_1_F2 population, individuals were classified as mutant when they exhibited more than 11% 4-seeded pods per plant. Based on this criterion, a discrete subset of plants was identified, and the segregation pattern was consistent with a 15:1 ratio (329 wild type: 20 mutant; χ² = 0.16, p = 0.69), supporting the involvement of two independently segregating recessive loci (Table S2).

We next examined the association between the CGH-defined deletion intervals and pod phenotype in BC_1_F4-derived lines. Lines carrying the chromosome 2 deletion (MO27-Chr.2) or the chromosome 6 deletion-containing interval (MO27-Chr.6) each produced a higher proportion of four-seeded pods than WT (Figure 2b, d). Progeny harboring both the chromosome 2 and chromosome 6 lesions (MO27-Chr.2/6) showed a further increase in four-seeded pod frequency, approximately two-fold higher than either single-deletion genotype (Figure 2b, d). Consistent with this shift in pod class distribution, average seeds per pod were significantly increased in the deletion lines, with the strongest effect observed in MO27-Chr.2/6 (Figure S2a). These data support a model in which the chromosome 2 and chromosome 6 intervals each contribute to the phenotype, with combined loss producing a stronger effect.

PCR-based genotyping confirmed the presence or absence of the chromosome 2 and/or chromosome 6 deletions in WT, MO27, and segregating progeny, supporting the genotype assignments used for phenotypic comparisons (Figure 2c,d). Although 100-seed weight was significantly but modestly reduced in the Chr.2/6 and Chr.6 deletion lines (Figure 2e), total pod number per plant and plant height were largely comparable among genotypes (Figure 2f, g). These results indicate that the primary developmental consequence of the Chr.2 and Chr.6 lesions is an alteration in seed number per pod, with a limited effect of the Chr.6 deletion on seed weight. Genetic annotation of the two co-segregating deletion intervals identified 43 predicted genes including *Glyma.02G202000*, encoding a predicted N-acyltransferase, within the chromosome 2 interval and a cluster of ULT-related genes within the chromosome 6 interval, including *Glyma.06G250300* and *Glyma.06G250500* (Table S1). Phylogenetic analysis supported the designation of *Glyma.02G202000* and its closest soybean paralog *Glyma.10G074700* as *GmNAT1* and *GmNAT2*, respectively, with the Arabidopsis and tomato homologs referred to as *AtNAT-like* (*AT3G52610*) and *SlNAT* (*Solyc09g009990*) (Figure S3a). A separate ULT phylogeny resolved soybean ULT-related genes into *GmULT*-*A1*–*A2* and *GmULT*-*B1*–*B4*, while two divergent sequences were retained as ULT-like genes (Figure S3b). Within the chromosome 6 interval, *GmULT*-*B1* and *GmULT*-*B3* were prioritized because both were deleted in MO27 and expressed in WT floral tissue (Figure S3c). qRT-PCR analysis of unopened floral buds showed that *GmNAT1* and *GmULT*-*B1* transcripts were detected in WT but absent or strongly reduced in MO27 and the corresponding single-deletion genotypes (Figure 2h,i), consistent with deletion-dependent disruption of candidate gene expression.

These results define candidate genes within the chromosome 2 (*i.e*., *GmNAT1*) and 6 (i.e., *GmULT*-*B1* and *GmULT*-*B3*) intervals and link their deletion to reduced gene expression in floral buds, providing a basis for further functional analysis.

### CRISPR/Cas9-mediated mutagenesis of *GmNAT1* and *GmULT* genes partially phenocopies the elevated four-seeded pod phenotype

To determine whether disruption of these candidates is sufficient to account for the elevated frequency of 4-SP observed in MO27, we generated targeted loss-of-function alleles by CRISPR/Cas9-mediated genome editing. A multiplexed CRISPR/Cas9 construct was assembled to simultaneously target *GmNAT1*, *GmULT-B1*, and *GmULT-B3* (Figure 3a). Transgenic soybean plants were produced by Agrobacterium-mediated transformation, and independent edited lines carrying mutations at the three loci were verified by Sanger sequencing (Figure S4).

**Figure 3.**
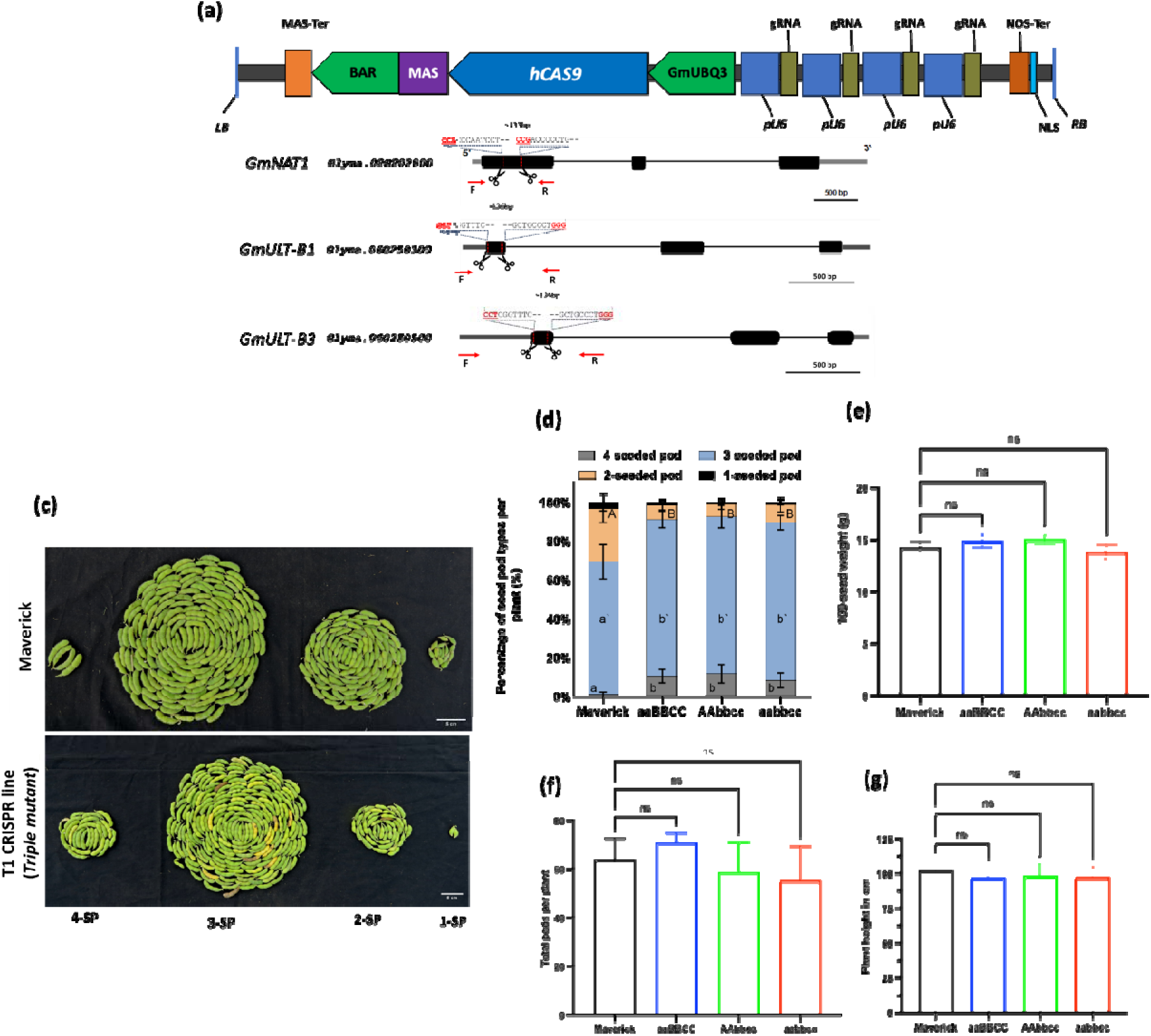
CRISPR/Cas9-mediated mutagenesis of GmNAT1 and GmULT validates thei contribution to four-seeded pod formation. (a) Schematic representation of the multiplex CRISPR/Cas9 construct carrying four guide RNAs targeting *GmNAT1* (*Glyma.02G202000*), *GmULT-B1* (*Glyma.06G250300*), and *GmULT-B3* (*Glyma.06G250500*), together with gene structures and corresponding CRISPR/Cas9 target sites. The stacked gRNA cassette was assembled in a soybean transformation vector containing BAR, MAS, hCas9, GmUBQ3, NLS, and NOS terminator elements. Protospacer adjacent motif sequences are shown in red, and arrows indicate PCR primers used for genotyping. Black boxe represent exons and lines represent introns or intergenic regions. Deleted regions generated by paired guide RNAs are indicated. (c) Representative pod sets from wild-type Maverick and the CRISPR/Cas9 T1 triple-mutant line 26-10, showing an increased proportion of four-seeded pods in the mutant. Pod categories are indicated below the images. Scale bars = 1 cm. (d) Percentage of four-seeded pods per plant in Maverick and CRISPR/Cas9 mutant genotypes. (e–g) Agronomic traits of wild-type and mutant plants, including 100-seed weight (e), total pod number per plant (f), and plant height (g). Data are presented as mean ± SE. Statistical significance was assessed by one-way ANOVA followed by Tukey’s multiple range test, with different letters indicating significant differences (P < 0.01). Uppercase letters denote wild-type alleles and lowercase letters denote CRISPR-induced mutant alleles at the GmNAT1 (A/a), GmULT-B1 (B/b), and GmULT-B3 (C/c) loci.

Phenotypic analysis of the resulting single and combination mutant lines demonstrated a significant increase in the proportion of 4-SP relative to wild-type plants (Figure 3c, d). The production of three-seeded pods (3-SPs) is also significantly increased in the CRISPR lines compared to wildtype, while that of two-seeded pods (2-SPs) decreased. Consistent with these changes in pod class distribution, the average number of seeds per pod was significantly increased in the mutant lines (Figure S2b). Thus, disruption of *GmNAT1* or *GmULT-B* , is sufficient to alter pod seed number distribution in the direction observed in MO27-Chr.2 and MO27-Chr.6, respectively. Notably, however, the magnitude of the increase in 4-SP in the triple mutant was consistently lower than that of the original FN mutant, indicating that the CRISPR-derived genotype reproduces the phenotype only in part rather than fully phenocopying MO27 (Figure 3d).

To assess whether this increase in seed number per pod was associated with broader pleiotropic effects on plant performance, we quantified 100-seed weight, total pod number per plant, and plant height. None of these traits differed significantly between CRISPR mutants and wild-type controls (Figure 3e–g). These data indicate that the increase in four-seeded pod frequency can occur without detectable penalties in the major agronomic traits evaluated under the conditions tested.

Taken together, these results provide direct genetic evidence that *GmNAT*, *GmULT*-*B1*, and *GmULT*-*B3* contribute functionally to the control of seed number per pod in soybean. At the same time, the inability of the triple mutant to fully recapitulate the MO27 phenotype strongly suggests that additional genetic factors within the FN deletion background, and/or higher-order combinatorial effects associated with the chromosome 2 and/or chromosome 6 lesions are required for full expression of the high four-seeded pod phenotype.

### Tomato ULT homologs modulate floral architecture and locule formation

To determine whether ULT-like genes may contribute to reproductive trait variation outside soybean, tomato was used as a heterologous validation system in which floral organ number and carpel-derived fruit traits can be readily quantified. Tomato ULT homologs were identified based on phylogenetic analysis using protein sequences from representative plant species. The resulting tree placed *SlULT1, SlULT2,* and *SlULT3* within the ULT-like lineage, providing a phylogenetic framework for cross-species functional analysis (Figure S3a, b).

To test their developmental roles directly, CRISPR/Cas9 knockout lines were generated in tomato cultivar M82 targeting three ULT-like genes, *SlULT1, SlULT2,* and *SlULT3*. Independent edited lines were recovered and molecularly validated by sequence analysis of the targeted loci, confirming mutagenesis at the expected positions (Figure S5a, b, c). Phenotypic analysis demonstrated that disruption of *SlULT1, SlULT2,* and *SlULT3* increased floral organ number relative to M82. The *slult* triple mutant showed increased numbers of sepals, petals, and stamens, together with altered floral architecture (Figure 4a, b).

**Figure 4.**
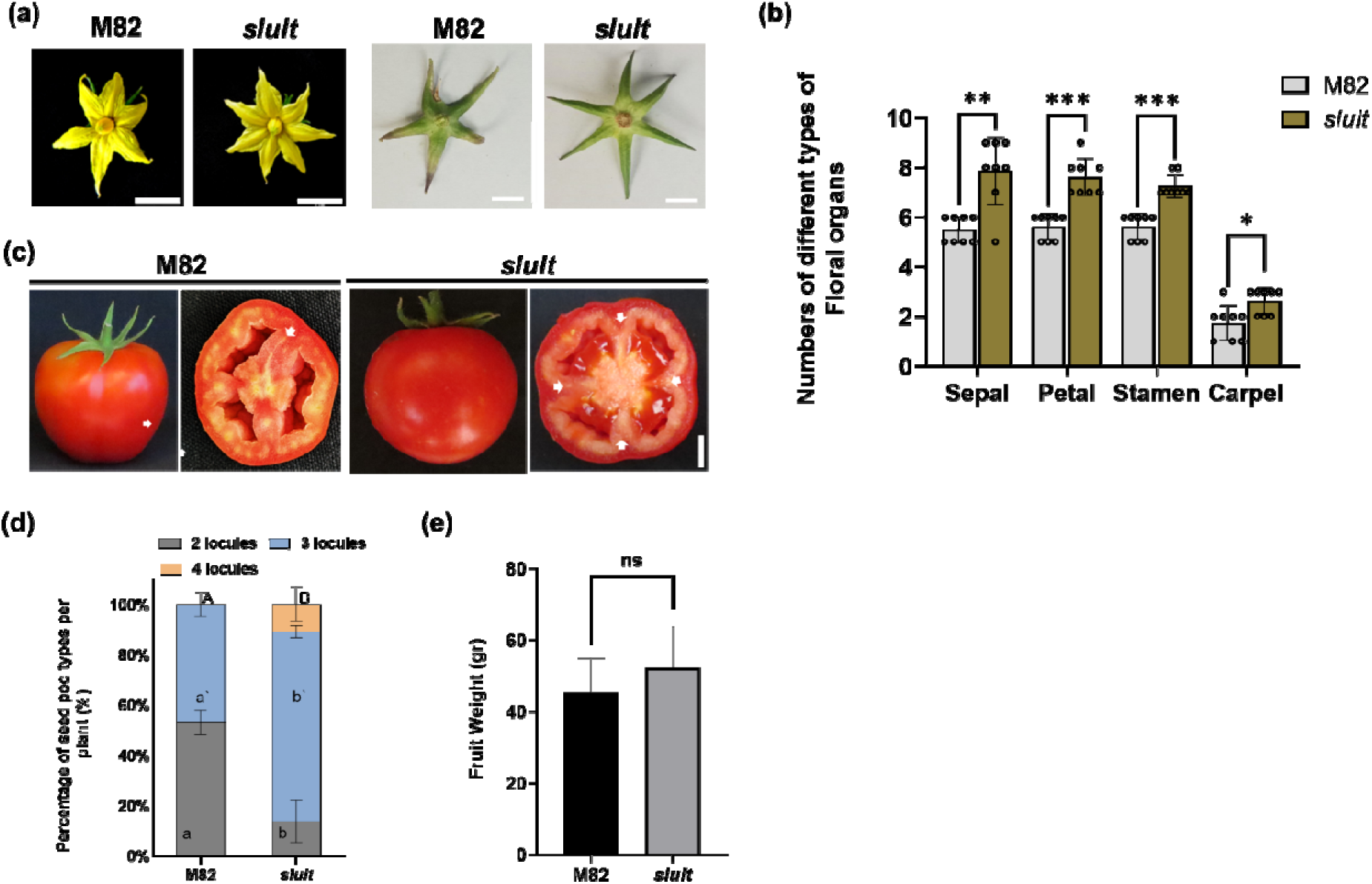
CRISPR/Cas9-engineered *slult* tomato mutants display increased floral organ number and enhanced fruit locule traits. (a) Representative frontal views of flowers and sepal from M82 and *slult* plants. Here, *slult* indicates the *slult1/2/3* triple mutant. Scale bar = 1 cm. (b) Quantification of floral organ number in WT and mutant flowers. Data are shown as mean ± SD n = 30. Asterisks indicate significant differences between genotypes as determined by unpaired two-tailed Welch’s t-test (*P < 0.05, **P < 0.01, ***P < 0.001). (c) Representative images of mature fruits and transverse sections. Scale bar = 1 cm. (d) Quantification of locule number in WT and mutant fruits. Data are shown as mean ± SD. Statistical significance was determined using an unpaired two-tailed Welch’s t-test. (e) Fruit weight in M82 and slult plants. Data are shown as mean ± SD (n ≥ 36 fruits per genotype). Statistical significance was determined using an unpaired two-tailed Welch’s t-test to account for unequal sample sizes between groups. (ns = not significant)

Consistent with the altered ovary morphology, mature fruits of *slult* plants showed increased locule number relative to M82 (Figure 4c, d). Fruit weight was not significantly different between M82 and *slult* plants, indicating that increased locule number in the *slult* mutant was not accompanied by a detectable increase in overall fruit mass under the conditions tested (Figure 4e).

Together, these results show that disruption of tomato ULT-like genes affects floral organ patterning and carpel-derived fruit morphology. These phenotypes are broadly consistent with the proposed contribution of ULT-like genes to reproductive organ development in soybean deletion lines, supporting the use of tomato as a heterologous system to test candidate reproductive regulators identified from the soybean deletion intervals.

### Interaction analysis reveals a Chr.6-dominant transcriptional state and module-specific non-additive regulation in the four-seeded pod mutant

Soybean CRISPR/Cas-derived NAT and ULT mutants exhibited increased four-seeded pod frequency, but failed to fully recapitulate the strong phenotype observed in the FN-MO27 mutant background. These observations support a role for the NAT–ULT module in seed number regulation, while also suggesting that the enhanced phenotype of the MO27 mutant likely reflects combinatorial effects involving additional genetic components located within the Chr.2 and/or Chr.6 deletion regions.

To investigate this possibility, we compared RNA-seq profiles from MO27-Chr.2, MO27-Chr.6, and MO27-Chr.2/6 during early floral development and performed interaction analysis using the DESeq2 interaction model. This framework separates the additive component expected when two genetic factors act independently from the interaction component that emerges when both factors coexist in the same genetic background. Accordingly, the analysis allowed us to determine whether the transcriptional state of MO27-Chr.2/6 could be explained by the additive combination of the Chr.2 and Chr.6 deletion effects, or whether specific gene groups exhibited non-additive responses resulting from combinatorial perturbation. Genes were subsequently categorized according to their expression patterns as Chr.2-driven, Chr.6-driven, buffered, complex_interaction, or double_specific. Chr.2-driven and Chr.6-driven genes were defined as genes whose expression patterns in MO27-Chr.2/6 largely resembled those observed in the corresponding single-deletion lines, whereas buffered and complex_interaction categories were used to interpret transcriptional responses deviating from additive expectations.

Interaction analysis revealed that the transcriptional architecture of MO27-Chr.2/6 was strongly dominated by the Chr.6 deletion effect. Among 599 classified genes, Chr.6-driven upregulated and Chr.6-driven downregulated genes comprised the largest categories, with 185 and 148 genes, respectively, together accounting for 55.6% of all classified genes (Figure 5a; Table S3). In contrast, Chr.2-driven genes were rare, consisting of only three upregulated and two downregulated genes, while double_specific genes were also limited in number. These results indicate that the transcriptional state of MO27-Chr.2/6 does not represent a broadly distinct double-mutant-specific program, but instead largely reflects a Chr.6-dominant transcriptional state resembling that of MO27-Chr.6.

**Figure 5.**
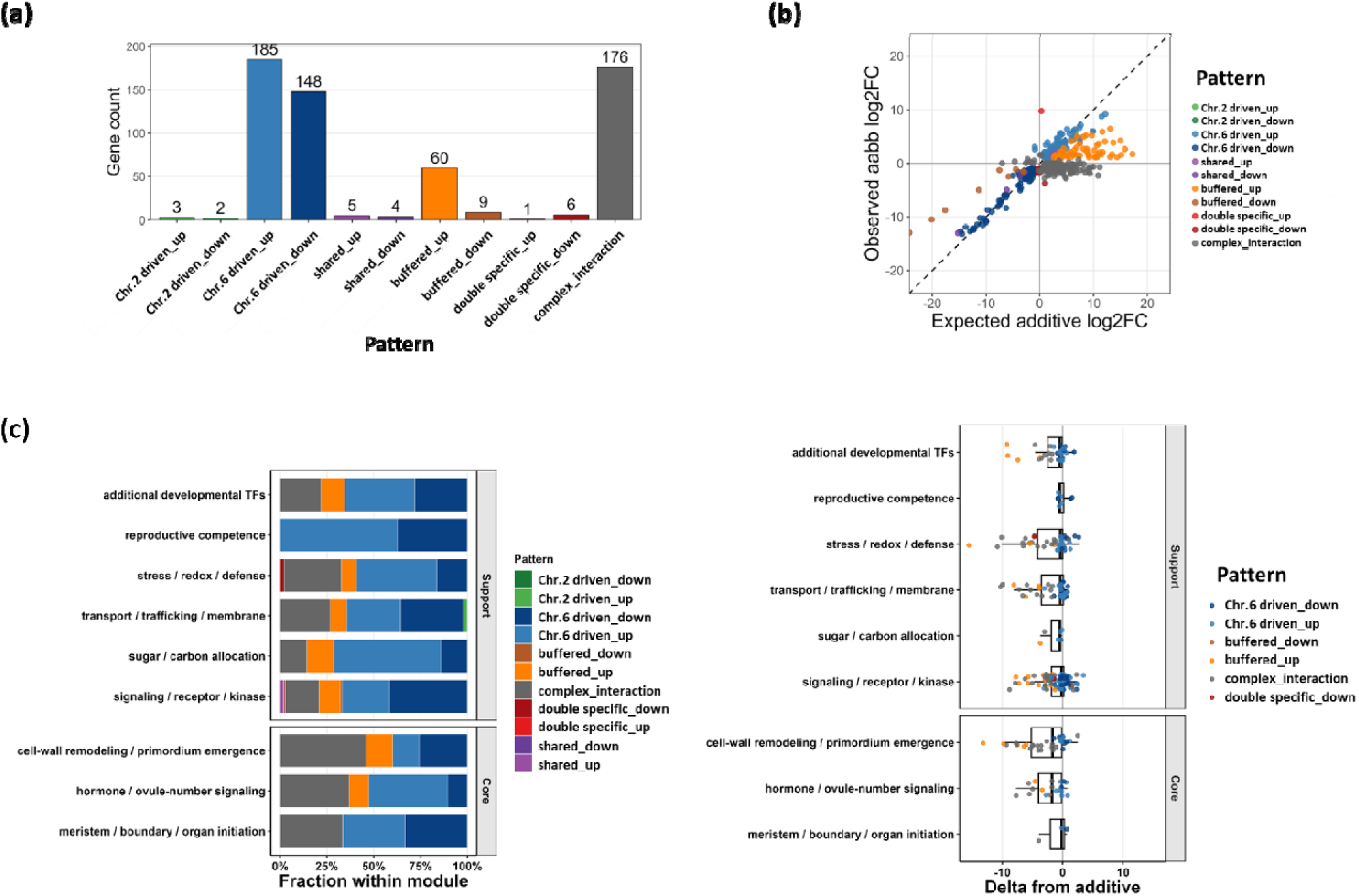
Interaction-driven transcriptional reprogramming underlying the four-seeded pod phenotype. (a) Distribution of genes across interaction-defined expression patterns. Genes were classified based on their deviation from the expected additive model derived from the two single-deletion lines, MO27-Chr.2 and MO27-Chr.6. Categories include Chr.2-driven, Chr.6-driven, shared, buffered, double-specific, and complex interaction patterns, stratified by up- and down-regulation. Gene counts for each category are indicated. (b) Comparison of observed versus expected additive log2 fold changes for each gene. The x-axis represents the expected additive effect calculated from MO27-Chr.2 and MO27-Chr.6, and the y-axis represents the observed expression change in MO27-Chr.2/6. The dashed diagonal indicates perfect additivity. Deviations from this line reflect non-additive genetic interactions, with points colored according to interaction-pattern classification. (c) Functional module-level distribution of interaction patterns. Genes were grouped into biologically defined modules, and the proportion of each interaction pattern within each module is shown. Core modules include meristem/boundary/organ initiation, hormone/ovule-number signaling, and cell wall remodeling/primordium emergence. Support modules include signaling/receptor/kinase, metabolism, transport/trafficking/membrane, stress response, reproductive competence, and additional developmental transcription factors. (d) Module-wise distribution of deviation from additivity. Each point represents an individual gene, and boxplots summarize the distribution within each functional module. This analysis highlights module-specific differences in the extent of additive deviation, including buffered and complex interaction behaviors.

Additive deviation analysis further supported this interpretation (Figure 5b). Most Chr.6-driven genes clustered near the additive diagonal, indicating that expression changes observed in MO27-Chr.2/6 were largely predictable from the Chr.6 deletion effect alone. In contrast, buffered genes exhibited attenuated responses relative to expectation, whereas complex_interaction genes deviated broadly from additive expectations, indicating non-linear combinatorial regulation. These results suggest that although the overall transcriptional landscape of MO27-Chr.2/6 is primarily governed by a Chr.6-dependent background, selective groups of genes undergo additional non-additive regulation resulting from the combined deletions.

To determine whether these transcriptional interaction patterns were functionally organized, genes were grouped into developmental and biological modules based on annotation and ortholog-informed functional classification. Several support-related categories, including signaling/receptor/kinase, transport/trafficking/membrane, and reproductive competence modules, were predominantly composed of Chr.6-driven genes (Figure 5c; Table S3). These findings indicate that the Chr.6 deletion broadly establishes the transcriptional framework of MO27-Chr.2/6 across multiple biological layers.

However, not all modules responded equivalently. The hormone/ovule-number signaling module and the cell wall remodeling/primordium emergence module contained relatively high proportions of buffered and complex_interaction genes. This indicates that these pathways are not simply passive outputs of the Chr.6 deletion effect, but are selectively reconfigured in the combined MO27-Chr.2/6 background. Because ovule number regulation, hormone signaling, and primordium emergence are directly associated with seed number determination, the non-additive responses observed within these modules likely represent key mechanistic components underlying the enhanced four-seeded pod phenotype of the FN mutant.

Delta-from-additivity analysis further reinforced this interpretation (Figure 5d). Most modules remained centered near zero, indicating broadly additive or Chr.6-dominant behavior across much of the transcriptome. However, the hormone/ovule-number signaling and cell wall remodeling/primordium emergence modules displayed substantially broader distributions and contained larger numbers of genes deviating from additive expectations. In contrast, reproductive competence and meristem/boundary/organ initiation modules exhibited comparatively narrow distributions, suggesting that these pathways maintain relatively stable or predictable transcriptional responses despite the strong developmental phenotype of MO27-Chr.2/6.

Notably, the cell wall remodeling/primordium emergence module exhibited the broadest distribution of non-additive responses. Ovule primordium emergence is tightly associated with boundary flexibility, cell wall loosening, cell wall reorganization, and primordium outgrowth. Therefore, the selective deviation from additivity observed in this module suggests that developmental processes occurring after ovule primordium initiation, including primordium emergence or stabilization, may be specifically reprogrammed in MO27-Chr.2/6. Similarly, the increased non-additive responses observed within the hormone/ovule-number signaling module support the possibility that signaling thresholds associated with ovule initiation or ovule number determination are reset in the double-deletion background.

Collectively, the interaction analysis demonstrates that the transcriptional state of the FN-derived MO27-Chr.2/6 mutant cannot be adequately explained by the simple additive combination of the two single-deletion effects or by genome-wide synergistic amplification. Instead, the data support a hierarchical model in which the Chr.6-associated deletion establishes the dominant transcriptional background, upon which selective non-additive reprogramming emerges within hormone/ovule-number signaling and cell wall remodeling/primordium emergence modules. This layered regulatory architecture provides a mechanistic framework for understanding how core NAT–ULT perturbation, a Chr.6-dominant transcriptional state, and module-specific non-additive developmental reprogramming collectively contribute to the enhanced four-seeded pod phenotype observed in the soybean FN mutant.

### NAT–ULT perturbation separates a conserved developmental output from FN background-dependent responses

To define the transcriptional basis of the four-seeded pod phenotype, we integrated RNA-seq profiles from FN deletion mutants with qRT–PCR validation in both FN-derived and CRISPR/Cas9-generated lines. We focused on gene groups associated with ovule initiation, meristem maintenance, reproductive developmental regulation, and cell wall remodeling to distinguish conserved NAT–ULT downstream outputs from responses requiring the broader FN deletion background.

We first examined TCP7/21-like genes because TCP transcription factors are linked to meristem boundary formation and lateral organ initiation. Among five soybean TCP7/21-like orthologs— *Glyma.10G285900*, *Glyma.09G284500*, *Glyma.09G284300*, *Glyma.20G001600*, and *Glyma.20G103400*—*Glyma.09G284300* was significantly upregulated in all FN deletion mutant genotypes (Figure 6a). Because this gene is highly similar to its duplicated paralog *Glyma.09G284500*, gene-specific qRT–PCR validation was not technically feasible. however, examined tomato orthologs and found that the expression of *Solyc.04G009180*, one of two tomato TCP7/21-like genes, was significantly upregulated in the *slult* triple mutant (Figure 6d). These results and the RNA-seq data in soybean, therefore, support TCP7/21-like activation as a conserved downstream response associated with NAT1 and ULT perturbation.

**Figure 6.**
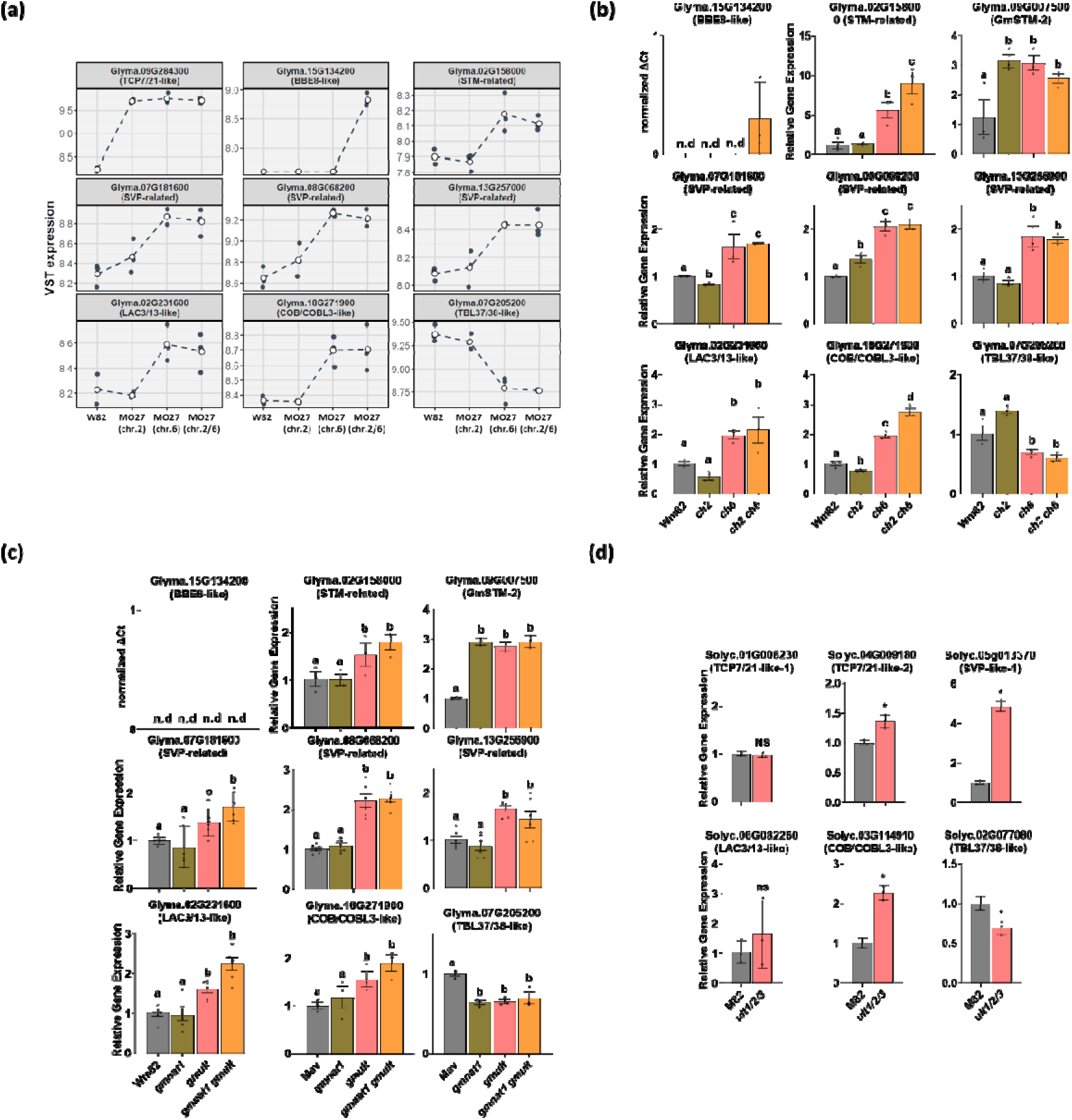
Representative gene expression profiles and experimental validation across genetic backgrounds and species. (a) Variance-stabilized (VST) expression profiles of selected representative genes across four genotypes (W82, MO27(chr.2), MO27(chr.6), MO27(chr.2/6)). Points indicate biological replicates, and dashed lines represent mean expression trends across genotypes. (b) qRT–PCR validation of selected genes in the FN mutant background. Expression patterns were assessed to confirm RNA-seq-derived trends, focusing on both common-output genes and genotype-specific responses. (c) qRT–PCR validation in CRISPR/Cas9-generated soybean mutants targeting NAT1 and ULTs loci. (d) qRT-PCR validation in CRISPR/Cas9-generated tomato mutant targeting ULTs loci. For qRT–PCR analyses, relative expression was calculated using the ΔΔCt method after normalization to the internal reference gene. Soybean samples were normalized to EF1, and tomato samples were normalized to TIP41. Data are shown as mean ± SD from biological replicates. For panels (b) and (c), statistical significance was determined by one-way ANOVA followed by Tukey’s multiple-comparison test, and different lowercase letters indicate statistically significant differences among genotypes (P < 0.05). For panel (d), statistical significance between WT and mutant samples was determined using an unpaired two-tailed Welch’s t-test. n.d. indicates not detected.

STM-related regulation was also reproducibly associated with the four-seeded pod phenotype. The STM-related gene *Glyma.02G158000* was significantly upregulated in chr.6 and chr.2/6 FN mutants by both RNA-seq and qRT–PCR, and was also induced in CRISPR/Cas9-generated *ult* and *nat1 ult* mutants (Figure 6a, b, c). Among four canonical soybean STM orthologs— *Glyma.07G263600* (*GmSTM-1*), *Glyma.09G007500* (*GmSTM-2*), *Glyma.15G111900* (*GmSTM-3*), and *Glyma.17G010200* (*GmSTM-4*)—*GmSTM-2* was significantly upregulated across all mutant genotypes in both FN and CRISPR backgrounds (Figure 6b, c). Thus, STM-related transcriptional activation, together with TCP7/21-like induction, represents a core regulatory output repeatedly associated with NAT–ULT perturbation.

Additional transcriptional responses showed stronger chr.6 deletion or *ult* dependency. SHORT VEGETATIVE PHASE (SVP)/AG-like MADS-box genes, which are associated with reproductive developmental regulation (Gregis *et al*., 2013; Mateos *et al*., 2015), and cell wall modification-associated genes were mainly altered in chr.6 deletion or *ult* backgrounds (Figure 6a, b, c). Similar SVP/AG-like induction was observed in tomato *ult* mutants, whereas LAC3/13-like genes did not show clear changes in tomato (Figure 6d). These results suggest that ULT perturbation activates a reproductive and cell wall-associated transcriptional branch that is only partially conserved across species.

Finally, *Glyma.15G134200* (*GmBBE8L*), an *AtBBE8-like* putatively linked to apoplastic redox activity and cell-wall modification, showed a distinct FN double-specific pattern. It was undetectable in WT, chr.2, and chr.6 single mutants but strongly induced in the chr.2/6 double deletion mutant, as confirmed by qRT–PCR (Figure 6a, b). In contrast, no detectable expression was observed in the CRISPR/Cas9 validation set, including wild-type, *gmnat1*, *gmult*, and *gmnat1 gmult* plants (Figure 6c). Thus, *GmBBE8L* is unlikely to represent a general downstream target of NAT–ULT disruption. Instead, its FN double-specific induction may reflect the combined loss of *GmNAT1*, *GmULT-B* genes, and additional co-deleted gene(s) on chromosome 2 and/or chromosome 6, which together establish a transcriptional state that is not reproduced in the CRISPR/Cas9 *gmnat1 gmult* mutants.

## Discussion

### A Hierarchical NAT–ULT Downstream Model for the Four-Seeded Pod Phenotype

The combined transcriptional and genetic analyses support a hierarchical model in which NAT– ULT perturbation generates multiple downstream outputs with distinct functional and genetic dependencies. Rather than acting through a single linear pathway, the four-seeded pod phenotype appears to arise through sequentially layered effects on developmental competence, reproductive patterning, cell wall remodeling, and FN background-specific reinforcement.

In this model, Layer I represents a conserved core developmental response. NAT–ULT perturbation induces TCP7/21-like and STM-related transcriptional activation across independent mutant systems and genetic backgrounds, suggesting that these genes represent a stable downstream output rather than stochastic transcriptional variation. Functionally, this layer is consistent with increased meristematic competence and enhanced ovule initiation potential. STM-related genes are required for meristem maintenance, carpel initiation, and reproductive meristematic tissue development, including placental tissues from which ovule primordia arise (Clark *et al*., 1996; Scofield *et al*., 2007). ULT1 and STM also act antagonistically within the same meristematic network, as *ult1* mutations can restore organ-forming capacity to stm null mutants, indicating that this relationship directly influences lateral organ formation (Carles *et al*., 2004).

TCP7/21-like transcription factors may further contribute to meristematic boundary formation and lateral organ initiation. TCP factors regulate reproductive developmental genes and have been linked to ovule initiation and primordium arrangement; altered CIN-clade TCP activity disrupts ovule development and patterning in Arabidopsis (Viola and Gonzalez, 2023). Thus, TCP7/21/STM-related activation likely establishes a permissive developmental state that increases the probability of ovule primordium initiation. However, because these genes were already induced in single mutants and did not necessarily show further enhancement in double mutants, this conserved core layer is unlikely to fully explain the stronger FN double-mutant synergy.

Layer II builds on this permissive state through ULT-dependent activation of SVP/AG-like MADS-box genes and cell wall modification-associated genes, indicating that ULT loss activates an additional branch coupling reproductive patterning with structural remodeling. AG-clade MADS-box proteins, including SEEDSTICK, SHATTERPROOF1/2, and AG, form complexes with SEPALLATA proteins to specify ovule identity and guide primordium morphogenesis (Favaro *et al*., 2003). BELL1 and AG also influence the balance between carpel identity and ovule identity at the placental surface (Western and Haughn, 1999), while BELL1–MADS interactions suppress WUS during ovule development to maintain proper primordial cell identity (Brambilla *et al*., 2007). Within this framework, ULT-dependent induction of SVP/AG-like genes can be interpreted as relief of ULT-mediated epigenetic repression at reproductive developmental loci (Monfared *et al*., 2013).

In parallel, the induction of LAC3/13-like, COB/COBL3-like, and TBL37/38-like genes suggests that the chromosome 6/ULT-dependent layer also activates a cell wall remodeling program associated with reproductive organ development and seed formation. Ovule primordium initiation and subsequent seed development require coordinated changes in cell wall architecture, including localized wall loosening, cellulose organization, pectin modification, and lignin deposition. Laccase-family proteins promote oxidative monolignol polymerization and spatially regulated lignification (Hoffmann *et al*., 2020; Schuetz *et al*., 2014), and LAC3/13-related genes have been implicated in seed coat lignin deposition, seed viability, germination, and seed-setting processes (Yu *et al*., 2017; Zhuang *et al*., 2025). COBRA and COBRA-LIKE proteins regulate cellulose microfibril orientation and deposition, with COBL family members showing tissue- and stage-specific functions in Arabidopsis and soybean (Brady *et al*., 2007; Roudier *et al*., 2005; Sangi *et al*., 2021). TBL-family proteins modify wall polysaccharide acetylation, including pectin-related cell wall properties, which may influence the mechanical environment required for ovule primordium emergence and seed development (Dauphin *et al*., 2024; Gille and Pauly, 2012; Zhong *et al*., 2024). Consistent with the proposed involvement of cell wall remodeling in the 4-SP trait, genes related to pectin modification have been identified as possible mediators of this trait in soybean (Chen *et al*., 2023; Fang *et al*., 2021).

Therefore, Layer II may not simply promote ovule initiation, but may also modify the mechanical environment in which newly initiated primordia are stabilized and expanded. Because some cell wall-associated responses were not conserved in tomato, this layer likely includes both conserved developmental outputs and species-specific regulatory adaptations.

Layer III is superimposed on the first two layers and is defined by FN double-specific activation of *GmBBE8L*. Unlike the TCP7/21/STM core module or the broader ULT-dependent branch, *GmBBE8L* was activated specifically in the FN chr.2/6 double mutant and not in CRISPR/Cas9-generated nat1 ult double mutants. This indicates that simultaneous disruption of NAT and ULT alone is insufficient to induce this response and that additional modifier loci within the FN deletion intervals are likely required. GmBBE8L therefore represents an FN background-dependent transcriptional effector associated with stronger penetrance of the four-seeded pod phenotype.

Functionally, GmBBE8L may connect FN-specific transcriptional responses with localized cell wall reinforcement during ovule development. Its placement within Group 5 of the BBE-like family suggests a closer relationship to plant-specific extracellular oxidoreductases involved in monolignol or phenylpropanoid-derived alcohol oxidation than to canonical berberine alkaloid metabolism. Consistent with this interpretation, related BBE-like proteins have been associated with apoplastic redox activity and cell wall-associated oxidative metabolism (Daniel *et al*., 2015). Although the biochemical activity of GmBBE8L remains untested, its induction specifically in the FN double mutant supports a model in which developmental competence and ULT-dependent boundary remodeling are accompanied by phenylpropanoid-linked oxidative wall reinforcement, potentially stabilizing nascent ovule primordia and contributing to the enhanced four-seeded pod phenotype.

Collectively, this model explains how a shared NAT–ULT developmental module can generate a basal four-seeded pod phenotype, while FN-specific modifier responses amplify the phenotype through additional cell wall-associated reinforcement mechanisms. TCP7/21/STM-related activation first establishes a permissive state for increased ovule initiation; ULT-dependent SVP/AG-like and cell wall-associated pathways then remodel the reproductive and structural environment surrounding nascent ovule primordia; and FN double-specific activation of GmBBE8L may further stabilize newly initiated primordia, increase developmental persistence, or reduce developmental abortion through oxidative wall remodeling or boundary reinforcement (Figure 7).

**Figure 7.**
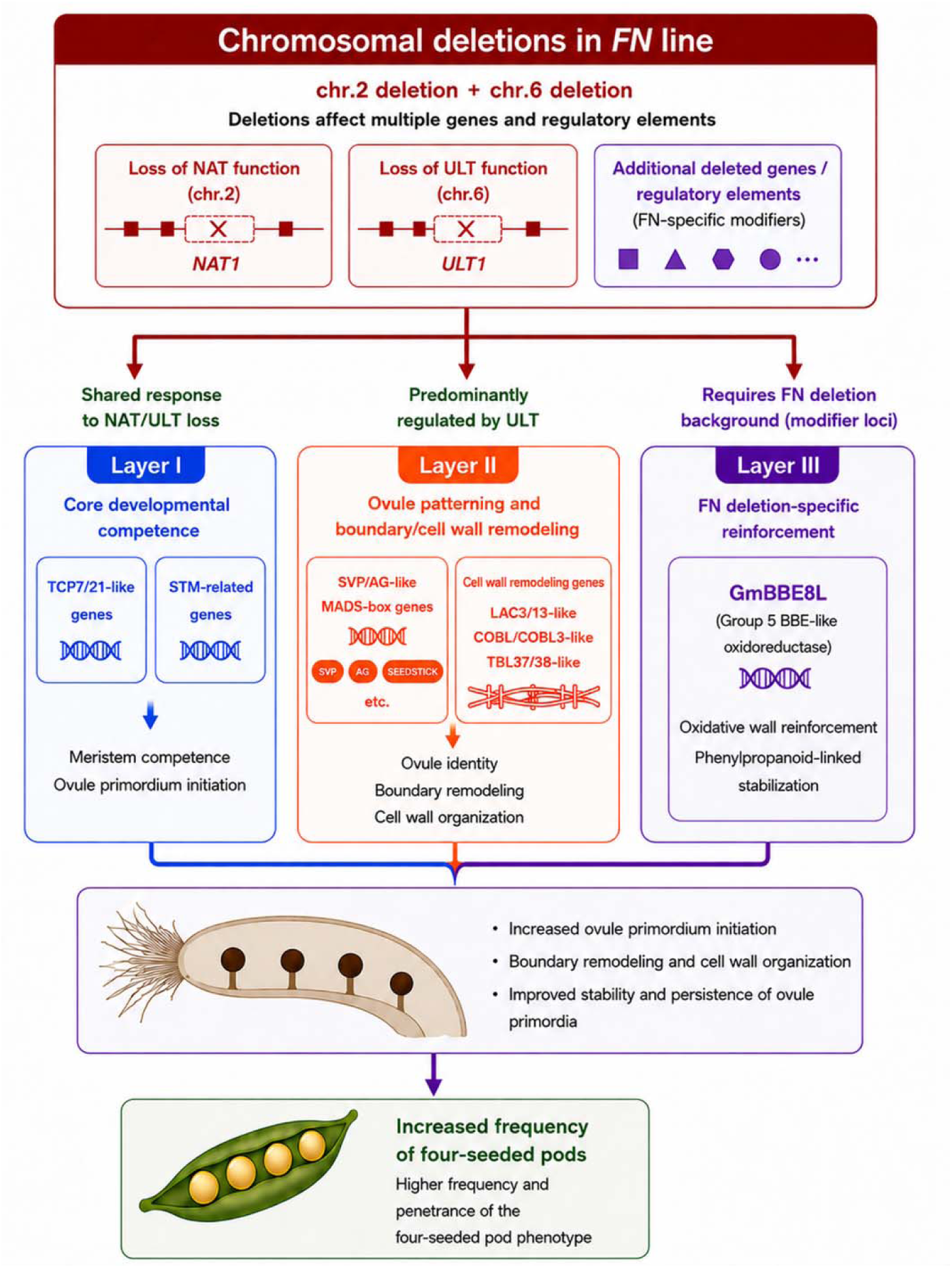
Parallel-layer model for chromosomal deletion-dependent regulation of soybean ovule development underlying the four-seeded pod phenotype. FN-induced deletions on chromosome 2 and chromosome 6 are proposed to affect multiple genes and regulatory elements, including loss of NAT and ULT function together with additional deletion-associated modifiers. These perturbations coordinately activate three parallel regulatory layers associated with enhanced ovule development. Layer I represents a conserved developmental competence module characterized by activation of TCP7/21-like and STM-related genes associated with meristem competence and ovule primordium initiation. Layer II represent reproductive patterning and cell wall remodeling pathways involving SVP/AG-like MADS-box genes together with LAC-, COBL-, and TBL-related cell wall regulators, potentially contributing to ovule identity establishment, boundary remodeling, and cell wall organization. Layer III represents an FN deletion-specific reinforcement module marked by induction of GmBBE8L, a Group 5 BBE-like oxidoreductase associated with oxidative cell wall reinforcement and phenylpropanoid-linked stabilization. The combined activities of these three layers are proposed to enhance ovule primordium initiation, remodeling, and persistence, ultimately increasing the frequency and penetrance of four-seeded pod formation in soybean.

This hierarchical framework provides a conceptual basis for a working model figure integrating developmental competence, reproductive patterning, and structural reinforcement as sequential regulatory layers downstream of NAT–ULT perturbation. Functional validation of GmBBE8L, together with analysis of responsive soybean COBL and LAC family members, is required to determine whether cell wall remodeling is a mechanistic driver rather than a correlative feature of the four-seeded pod phenotype.

### Species-Specific Differences in ULT-Related Reproductive Phenotypes: Soybean versus Tomato

Cross-species validation in tomato provided independent support for a conserved role of ULT-like genes in reproductive organ development. In tomato, CRISPR/Cas9-mediated disruption of *SlULT1*, *SlULT2*, and *SlULT3* increased floral organ number and altered floral architecture and ovary morphology. Mature fruits of *slult* plants also showed increased locule number relative to M82, although fruit weight was not significantly changed under the conditions tested. These results indicate that tomato ULT-like genes contribute to floral organ patterning and carpel-derived fruit morphology, consistent with the proposed involvement of soybean ULT-related genes in ovule and seed-number regulation.

However, the tomato phenotype should be interpreted as heterologous functional support rather than a direct phenocopy of the soybean deletion mutant. In soybean, the FN chromosome 6 deletion removes *GmULT-B1* and *GmULT-B2* and is associated with increased four-seeded pod frequency, particularly in combination with the chromosome 2 deletion. By contrast, the tomato experiment targeted three ULT-like genes in a species where floral organ number and locule number provide more tractable readouts of carpel and ovary patterning. Therefore, the shared feature between the two systems is not identical organ-level morphology, but rather the broader developmental sensitivity of reproductive meristems and carpel-derived structures to reduced ULT-like gene function.

One possible explanation for the difference in phenotypic severity and organ specificity is the distinct genomic context of the ULT gene family in soybean and tomato. Soybean has undergone whole-genome duplication, resulting in extensive gene family expansion and functional redundancy. In our phylogenetic analysis, soybean ULT-related genes were resolved into two major clades, designated *GmULT-A* (*GmULT-A1*–*A2*) and *GmULT-B* (*GmULT*-*B1*–*B4*). The FN deletion on chromosome 6 removes *GmULT*-*B1* and *GmULT*-*B2* but does not disrupt *GmULT-A* members, which may retain partial ULT pathway activity. This residual activity could buffer meristem expansion and restrict the phenotypic output primarily to quantitative changes in ovule or seed number rather than gross disruption of floral architecture.

In tomato, simultaneous disruption of *SlULT1*, *SlULT2*, and *SlULT3* likely reduces a larger fraction of total ULT-like activity, leading to a more visible effect on floral organ number and locule formation. This interpretation is consistent with the established role of *ULT1*-related activity in limiting meristematic potential through regulation of floral meristem programs, including *WUS*-associated stem cell maintenance. Because changes in meristem size and determinacy can alter carpel number and locule formation, the increased locule number observed in *slult* tomato fruits is compatible with reduced ULT-mediated restriction of reproductive meristem activity.

Taken together, the tomato data support the conclusion that ULT-like genes are conserved regulators of reproductive organ patterning, while the soybean data indicate that partial loss of a subset of GmULT genes can modify ovule or seed number without strongly perturbing overall pod or floral morphology. From an agronomic perspective, this distinction is important: partial redundancy within the soybean GmULT family may allow increased seed number per pod while avoiding the developmental penalties that could accompany stronger disruption of floral meristem regulation.

### Agricultural Significance and Prospects for Crop Improvement

Soybean is one of the most important global protein and oil crops, and increasing yield per unit area remains a central goal of breeding programs worldwide. Seed yield in soybean is a multiplicative function of plant number per area, pod number per plant, seed number per pod, and individual seed weight. While considerable progress has been made in maximizing pod number and seed fill efficiency, relatively little heritable variation in seed number per pod has been exploited in elite germplasm, largely because naturally occurring alleles that increase pod seed number are rare and the underlying genetic architecture was poorly understood. In contrast to recent QTL, GWAS, and QTN studies that provide genetic loci, candidate genes, and marker-assisted selection models for SNPP/NFSP, the NAT–ULT module provides a genetically validated entry point into the developmental control of ovule number itself (Li *et al*., 2021; Wang *et al*., 2024; Yang *et al*., 2013; Yuan *et al*., 2025).

The mutant germplasm described here—specifically, lines combining elevated four-seeded pod frequency with minimal change in individual seed weight—represents a class of agronomic resource that has not previously been well characterized in soybean. The ovule number per ovary sets the upper limit for seed number per pod, and because ovule number is established before fertilization, this trait is largely independent of post-fertilization environmental variation (Qadir *et al*., 2021). This developmental autonomy means that the seed number advantage should be stably expressed across diverse environments, a key requirement for agronomic utility. The finding that neither total pod number nor plant height was significantly altered in our mutants suggests that increased seed number per pod was not accompanied by obvious changes in these measured traits. However, because soybean yield components can compensate for one another, field-based evaluation of seed number per unit area, seed weight, and final yield will be required to determine whether this component-level change translates into yield gain (Vogel *et al*., 2021). The CLV-WUS signaling pathway and related meristem maintenance genes have been recognized as key targets for yield optimization across multiple crop species, including tomato, maize, and oilseed Brassica (Fletcher, 2018), and our demonstration that the NAT–ULT module functions as an upstream regulator of reproductive organ number adds a new layer of tractable molecular targets to this framework.

From a practical breeding perspective, the CRISPR/Cas9-derived triple mutant offers a particularly attractive platform because it achieves a significant increase in four-seeded pod frequency without any detectable change in 100-seed weight, total pod number, or plant height in the cultivar Maverick. The absence of pleiotropic penalties facilitates introgression into elite backgrounds without yield drag, and the defined molecular lesions in *GmNAT1*, *GmULT-B1*, and *GmULT-B2* can be readily tracked by molecular markers. Furthermore, the discovery that the FN background extends the phenotype beyond the CRISPR baseline suggests that additional gains may be achievable by targeting the NAC- and SPL-containing regulatory layer identified in our transcriptomic analysis, either by stacking additional CRISPR edits or by identifying natural allelic variation at these loci in germplasm collections. In conclusion, our study identifies a conserved NAT–ULT reproductive patterning module as a determinant of seed number per pod in soybean and shows that coordinated module-level disruption can increase four-seeded pod frequency without detectable reduction in seed weight in the CRISPR/Cas9-derived lines examined here. These findings provide mutant germplasm and candidate alleles for testing seed-number architecture as a yield-component trait in soybean breeding, while future field studies will be required to determine its impact on crop productivity (Vogel *et al*., 2021). These findings establish a mechanistic foundation for translating knowledge of floral organ patterning into practical strategies for increasing the productivity of one of the world’s most important crops.

### Experimental Procedures

#### Plant material, phenotyping, and growth conditions

The fast neutron-induced soybean (*Glycine max* [L.] Merr.) mutant MO27 was identified from a mutagenized Williams 82 (Wm82) population generated as previously described (Stacey et al., 2016). Initial phenotypic screening for altered seed number per pod was conducted using M plants. For genetic analysis, MO27 was backcrossed to Wm82 by manually pollinating emasculated Wm82 flowers with MO27 pollen. Backcross-derived populations and fixed progeny lines, including BC F to BC F generations, were cultivated under field conditions at the Bradford Research Center, University of Missouri, Columbia, during the 2018 and 2019 growing seasons.

CRISPR/Cas9-edited soybean mutant lines and tomato wild-type M82 and CRISPR/Cas9-edited *slult* mutant lines were maintained under greenhouse conditions. Phenotypic analyses included pod-type distribution, four-seeded pod frequency, total pod number per plant, plant height, seed morphology, floral organ number, fruit locule number, 100-seed weight, and fruit weight, depending on the experimental system analyzed.

#### Ovule number analysis

To determine whether increased seed number per pod originated prior to fertilization, unopened floral buds were collected and fixed in 70% (v/v) ethanol. Following removal of sepals and petals, ovaries were cleared in 30% (v/v) sodium hypochlorite solution (Clorox®) for 24 h with gentle rotation until tissues became translucent. Cleared ovaries were examined using a DM 5500B compound microscope (Leica Microsystems), and ovule numbers were recorded. Ovary and ovule dimensions were measured using ImageJ software.

#### Comparative genome hybridization and deletion genotyping

Comparative genome hybridization (CGH) was performed using a 696,139-feature soybean microarray platform as previously described (Bolon et al., 2011). Microarray probes were designed by Roche NimbleGen based on the Wm82 reference genome. Genomic DNA isolated from young leaves of MO27 and Wm82 using the DNeasy Plant Mini Kit (Qiagen) was labeled with Cy3 and Cy5 dyes, respectively. Hybridization, washing, scanning, and signal acquisition were performed according to the manufacturer’s instructions. Copy number variation events were identified using established log ratio thresholds to detect deletion and amplification intervals.

Candidate deletion loci were validated by PCR-based genotyping using primers spanning chromosome 2 and chromosome 6 deletion junctions. Co-segregation analyses between deletion genotypes and four-seeded pod frequency were subsequently performed using backcross-derived progeny populations.

#### RNA isolation and quantitative RT-PCR analysis

Total RNA was extracted from unopened floral buds using TRIzol reagent (Invitrogen) followed by purification with the RNeasy Plant Mini Kit (Qiagen). RNA samples were treated with TURBO DNase (Invitrogen) to remove contaminating genomic DNA. First-strand cDNA synthesis was performed using M-MLV reverse transcriptase (Promega).

Quantitative real-time PCR (qRT-PCR) was performed using SYBR Green chemistry on a QuantStudio real-time PCR system (Applied Biosystems). Soybean transcript levels were normalized against *EF1*, whereas tomato transcript levels were normalized against *TIP41*, which has been validated previously as a stable tomato reference gene (Expósito-Rodríguez *et al*., 2008). Relative transcript abundance was calculated using the ΔΔCt method. Primer sequences used for qRT-PCR are listed in Table S4.

#### CRISPR/Cas9 vector construction and plant transformation

CRISPR/Cas9 constructs targeting *GmNAT1* (Glyma.02G202000), *GmULT-B1* (Glyma.06G250300), and *GmULT-B3* (Glyma.06G250500) were generated for soybean genome editing. Single-guide RNA (sgRNA) target sequences were designed using the CCTop platform (Stemmer et al., 2015), with target sites selected within early coding exons to maximize loss-of-function mutagenesis.

Complementary oligonucleotides containing sgRNA sequences were synthesized, annealed, and cloned into the BbsI sites of pAtU6-26-SK to generate sgRNA expression cassettes. Correct assembly was confirmed by Sanger sequencing. Multiplex CRISPR/Cas9 constructs were assembled by inserting sgRNA expression modules together with a 35S-driven Cas9 cassette into the binary vector pFGC5941 (GenBank accession no. AY310901), following the dual CRISPR/Cas9 strategy described by Do et al. (2019). Binary vectors were introduced into *Agrobacterium tumefaciens* strain AGL1 by electroporation and subsequently used for soybean transformation.

For tomato genome editing, CRISPR/Cas9 constructs targeting *SlULT1*, *SlULT2*, and *SlULT3* were generated in the M82 background using the same strategy. sgRNA expression cassettes and Cas9 modules were assembled into pFGC5941 and introduced into *A. tumefaciens* strain AGL1 for tomato transformation.

#### Genotyping of CRISPR/Cas9-edited plants

Genomic DNA regions flanking CRISPR/Cas9 target sites were amplified using Phusion High-Fidelity DNA Polymerase and gene-specific primers listed in Table S4. PCR products were separated on 1.5% (w/v) agarose gels and purified either directly or following gel excision. Mutation events were confirmed by Sanger sequencing. Sequence alignments were performed using MUSCLE to identify insertions, deletions, and frameshift mutations. Stable inheritance of edited alleles was confirmed in subsequent generations prior to phenotypic characterization.

#### RNA sequencing and transcriptome analysis

RNA sequencing was performed using unopened floral buds collected from Wm82 and FN-derived genotypes carrying chromosome 2 deletions, chromosome 6 deletions, or combined chromosome 2/chromosome 6 deletions. Total RNA was extracted using TRIzol reagent and purified using the RNeasy Plant Mini Kit.

Poly(A)-selected RNA libraries were constructed using the Illumina TruSeq RNA Sample Preparation Kit v2 according to the manufacturer’s protocol. Library quality and concentration were evaluated using the KAPA Library Quantification Kit and a Caliper LabChip GX system (Caliper Life Sciences). Raw sequencing read quality was assessed using FastQC, and adapter trimming and low-quality base removal were performed using Trim Galore.

Processed reads were aligned to the *Glycine max* Wm82.a6.v1 reference genome and corresponding annotation files obtained from Phytozome using RNA STAR implemented through the Galaxy platform. Gene-level raw count matrices were subsequently generated using featureCounts.

Differential expression and interaction analyses were performed in R using DESeq2. An interaction model incorporating chromosome 2 deletion status, chromosome 6 deletion status, and their interaction term was used to identify additive and non-additive transcriptional responses associated with the combined deletion background. Genes were subsequently assigned to interaction-defined transcriptional classes, including chromosome 2-driven, chromosome 6-driven, buffered, complex interaction, and double-specific categories.

#### Phylogenetic analysis

Homologous NAT-like and ULT-like protein sequences were identified through BLAST searches against Phytozome and NCBI databases. Protein sequences from soybean, tomato, *Arabidopsis*, and representative plant species were aligned and analyzed using the NGPhylogeny.fr platform (<u>NGPhylogeny.fr</u>). Phylogenetic trees were visualized and annotated using iTOL (<u>Interactive</u> <u>Tree Of Life</u>).

### Statistical analysis

Statistical analyses were performed using Prism v10 (GraphPad Software) and R software where appropriate. For pairwise comparisons, Welch’s two-tailed t-test was applied when variances or sample sizes differed between groups. Multiple-group comparisons were performed using one-way ANOVA followed by Tukey’s multiple-comparison test or Welch’s one-way ANOVA followed by Games–Howell multiple-comparison test, as indicated in the corresponding figure legends. Chi-square tests were used for segregation analyses. Statistical significance was defined as P < 0.05 unless otherwise specified. Different lowercase letters were used to indicate statistically significant differences among groups; groups sharing the same letter were not significantly different.

---

**Supplementary Figure S1.**
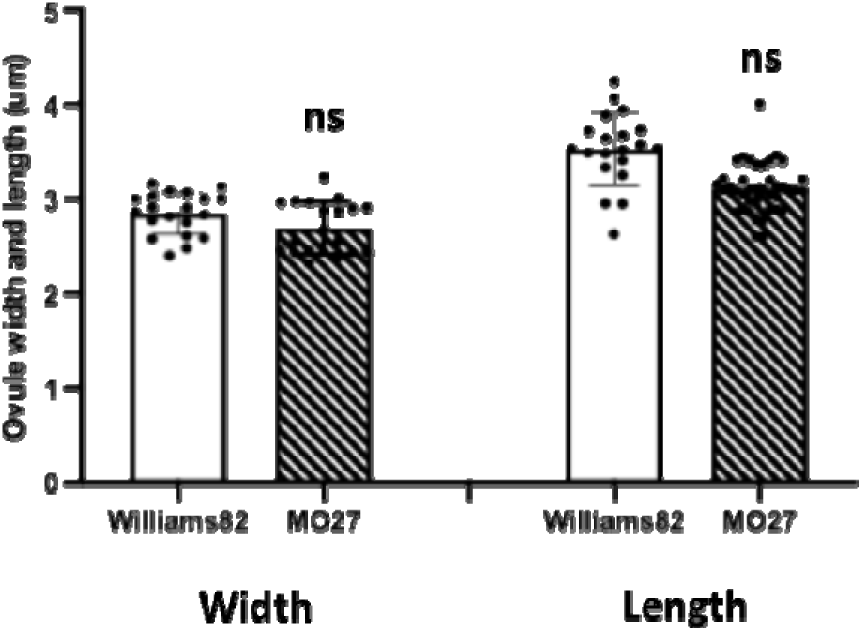
Morphometric analysis of ovules in WT and MO27. Quantification of ovule length and width in WT and MO27, showing no significant difference between genotypes (n = 50 ovaries). Data represent mean ± SD. Statistical analysis wa performed using Student’s *t*-test (P > 0.01).

**Supplementary Figure S2.**
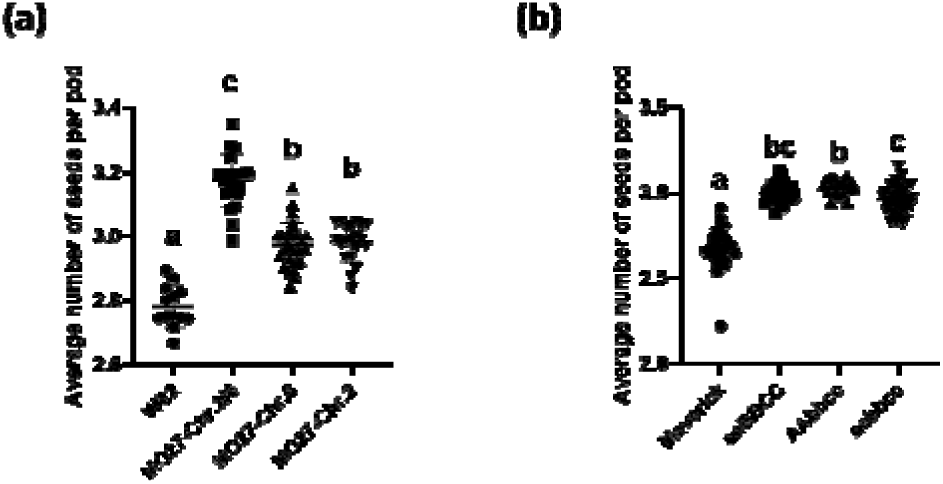
Increased average seed number per pod in FN deletion and CRISPR/Cas9-derived mutant lines. (a) Average seeds per pod in W82 and FN deletion mutants. (b) Average seeds per pod in Maverick and CRISPR/Cas9-derived mutants. Data are mean ± SD. Different lowercase letter indicate significant differences among genotypes as determined by Welch’s one-way ANOVA followed by Games–Howell multiple comparisons test (P < 0.05). Genotypes sharing a letter are not significantly different.

**Supplementary Figure S3.**
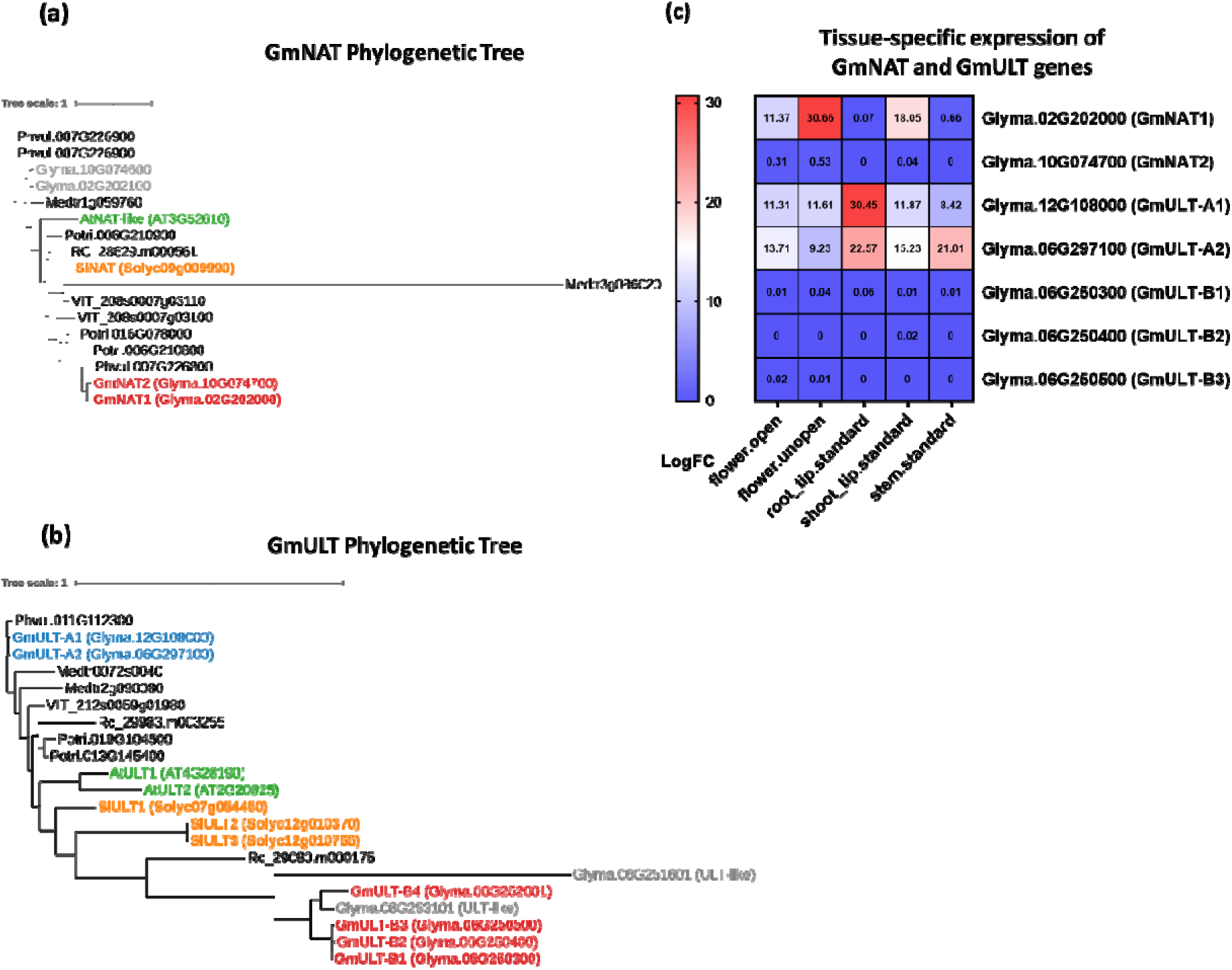
Phylogenetic identification and tissue expression of NAT and ULT genes. (a–b) Phylogenetic trees of NAT and ULT-related proteins from Glycine max, Arabidopsis thaliana, Solanum lycopersicum, Ricinus communis, Populus trichocarpa, and Vitis vinifera. Soybean genes were designated GmNAT1–GmNAT2 and GmULT-A1–A2/B1–B4 based on their evolutionary relationships with Arabidopsis NAT-like proteins and ULT1/2 homologs. (c) Tissue-specific expression profiles of selected soybean NAT and ULT genes in open flower, unopened flower, root tip, shoot tip, and stem. Nomalized expression values were derived from the JGI Plant Gene Atlas resource.

**Supplementary Figure S4.**
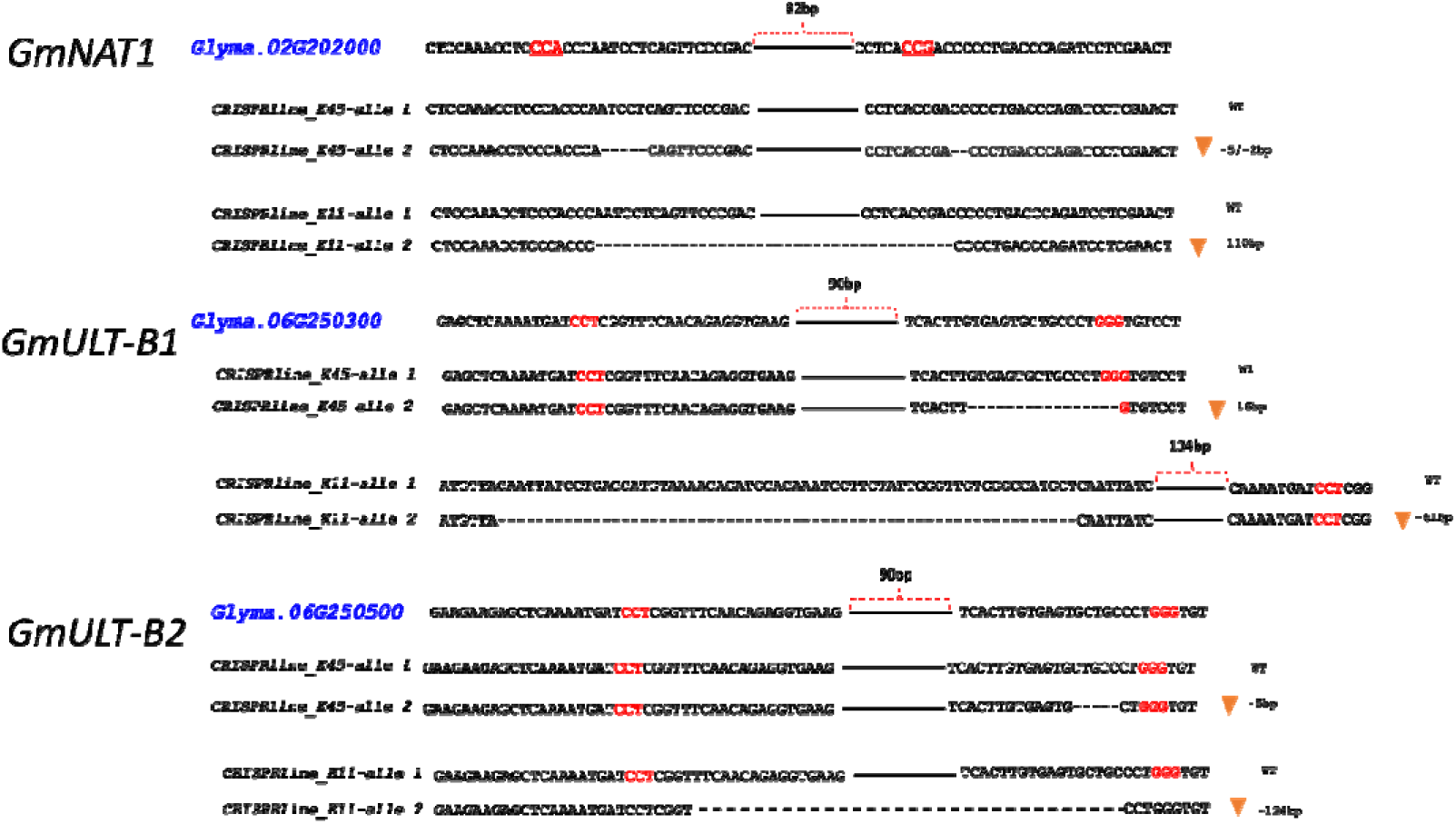
Sanger sequencing validation of CRISPR/Cas9-induced mutations in soybean GmNAT1, GmULT-B1, and GmULT-B3 alleles. Sequenced amplicons from CRISPR–WT lines matched the reference genome, whereas mutant lines carried deletions or indels at the expected target sites. PAM sequences are shown in red, deleted regions are indicated by brackets, and arrowheads mark the mutation positions.

**Supplementary Figure S5.**
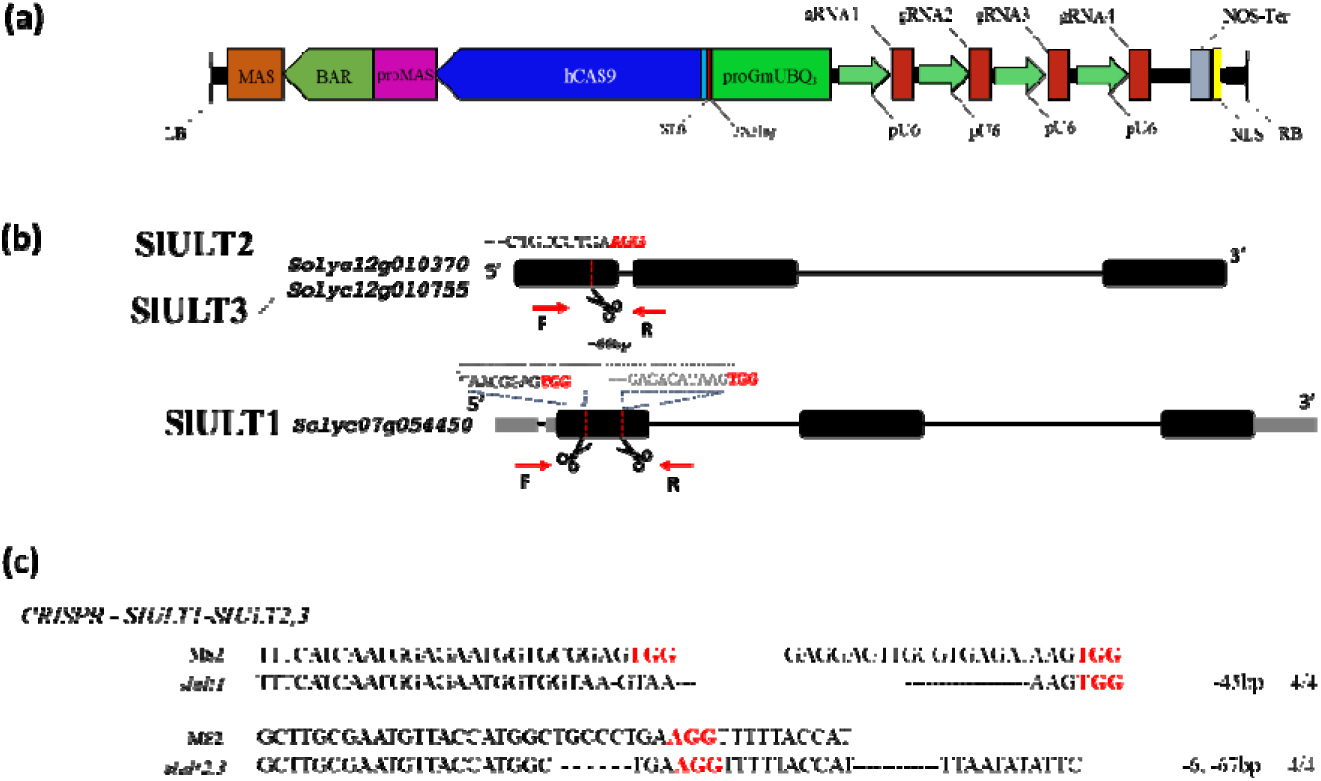
CRISPR/Cas9 construct design and target validation for ULT editing in tomato. (a) Schematic representation of the multiplex CRISPR/Cas9 construct used to target SlULT1, 2 and 3. The construct contains tandem U6 promoter-driven sgRNA cassettes assembled in a stacked array. Additional vector components include BAR, hCAS9, NLS, MAS, NOS terminator, and the soybean GmUBQ3 promoter. (b) Positions of the target sites within the SlULT1, 2 and 3 genomic regions. Black boxes indicate exons and lines indicate introns or intergenic regions. Red letters indicate PAM sequences, and red arrows mark the positions of gene-specific primers used for PCR genotyping. (c) Sequence alignments of wild-type M82 and representative CRISPR-edited alleles for SlULT1, 2 and 3. Deleted nucleotides are indicated by dashes, and PAM sequences are highlighted in red. The detected mutations are consistent with CRISPR/Cas9-mediated editing at the expected target sites.

## References

Ainsworth, E.A., Yendrek, C.R., Skoneczka, J.A., and Long, S.P. (2012) Accelerating yield potential in soybean: potential targets for biotechnological improvement. Plant, Cell & Environment, 35, 38–52.

Bolon, Y.-T., Haun, W.J., Xu, W.W., Grant, D., Stacey, M.G., Nelson, R.T., et al. (2011) Phenotypic and genomic analyses of a fast neutron mutant population resource in soybean. Plant Physiol, 156, 240–253.

Brady, S.M., Song, S., Dhugga, K.S., Rafalski, J.A., and Benfey, P.N. (2007) Combining Expression and Comparative Evolutionary Analysis. The COBRA Gene Family. Plant Physiol, 143, 172–187.

Brambilla, V., Battaglia, R., Colombo, M., Masiero, S., Bencivenga, S., Kater, M.M., and Colombo, L. (2007) Genetic and Molecular Interactions between BELL1 and MADS Box Factors Support Ovule Development in Arabidopsis. Plant Cell, 19, 2544–2556.

Carles, C.C., Choffnes-Inada, D., Reville, K., Lertpiriyapong, K., and Fletcher, J.C. (2005) ULTRAPETALA1 encodes a SAND domain putative transcriptional regulator that controls shoot and floral meristem activity in Arabidopsis. Development, 132, 897–911.

Carles, C.C. and Fletcher, J.C. (2009) The SAND domain protein ULTRAPETALA1 acts as a trithorax group factor to regulate cell fate in plants. Genes Dev., 23, 2723–2728.

Carles, C.C., Lertpiriyapong, K., Reville, K., and Fletcher, J.C. (2004) The ULTRAPETALA1 Gene Functions Early in Arabidopsis Development to Restrict Shoot Apical Meristem Activity and Acts Through WUSCHEL to Regulate Floral Meristem Determinacy. Genetics, 167, 1893–1903.

Chen, Y., Xiong, Y., Hong, H., Li, G., Gao, J., Guo, Q., et al. (2023) Genetic dissection of and genomic selection for seed weight, pod length, and pod width in soybean. The Crop Journal, 11, 832–841.

Chu, Y.-H., Jang, J.-C., Huang, Z., and van der Knaap, E. (2019) Tomato locule number and fruit size controlled by natural alleles of lc and fas. Plant Direct, 3, e00142.

Clark, S.E., Jacobsen, S.E., Levin, J.Z., and Meyerowitz, E.M. (1996) The CLAVATA and SHOOT MERISTEMLESS loci competitively regulate meristem activity in Arabidopsis. Development, 122, 1567–1575.

Cubas, P., Lauter, N., Doebley, J., and Coen, E. (1999) The TCP domain: a motif found in proteins regulating plant growth and development. The Plant Journal, 18, 215–222.

Cucinotta, M., Colombo, L., and Roig-Villanova, I. (2014) Ovule development, a new model for lateral organ formation. Front. Plant Sci., 5.

Cucinotta, M., Di Marzo, M., Guazzotti, A., De Folter, S., Kater, M.M., and Colombo, L. (2020) Gynoecium size and ovule number are interconnected traits that impact seed yield. Journal of Experimental Botany, 71, 2479–2489.

Daniel, B., Pavkov-Keller, T., Steiner, B., Dordic, A., Gutmann, A., Nidetzky, B., et al. (2015) Oxidation of Monolignols by Members of the Berberine Bridge Enzyme Family Suggests a Role in Plant Cell Wall Metabolism*. Journal of Biological Chemistry, 290, 18770–18781.

Dauphin, B.G., Ropartz, D., Ranocha, P., Rouffle, M., Carton, C., Le Ru, A., et al. (2024) TBL38 atypical homogalacturonan-acetylesterase activity and cell wall microdomain localization in Arabidopsis seed mucilage secretory cells. iScience, 27, 109666.

Expósito-Rodríguez, M., Borges, A.A., Borges-Pérez, A., and Pérez, J.A. (2008) Selection of internal control genes for quantitative real-time RT-PCR studies during tomato development process. BMC Plant Biol, 8, 131.

Fang, T., Bai, Y., Huang, W., Wu, Y., Yuan, Z., Luan, X., et al. (2021) Identification of Potential Gene Regulatory Pathways Affecting the Ratio of Four-Seed Pod in Soybean. Front. Genet., 12.

Favaro, R., Pinyopich, A., Battaglia, R., Kooiker, M., Borghi, L., Ditta, G., et al. (2003) MADS-Box Protein Complexes Control Carpel and Ovule Development in Arabidopsis. Plant Cell, 15, 2603–2611.

Fletcher, J.C. (2018) The CLV-WUS Stem Cell Signaling Pathway: A Roadmap to Crop Yield Optimization. Plants, 7.

Fletcher, J.C. (2001) The ULTRAPETALA gene controls shoot and floral meristem size in Arabidopsis. Development, 128, 1323–1333.

Galbiati, F., Sinha Roy, D., Simonini, S., Cucinotta, M., Ceccato, L., Cuesta, C., et al. (2013) An integrative model of the control of ovule primordia formation. The Plant Journal, 76, 446–455.

Gille, S. and Pauly, M. (2012) O-Acetylation of Plant Cell Wall Polysaccharides. Front. Plant Sci., 3.

Gregis, V., Andrés, F., Sessa, A., Guerra, R.F., Simonini, S., Mateos, J.L., et al. (2013) Identification of pathways directly regulated by SHORT VEGETATIVE PHASE during vegetative and reproductive development in Arabidopsis. Genome Biol, 14, R56.

Hoffmann, N., Benske, A., Betz, H., Schuetz, M., and Samuels, A.L. (2020) Laccases and Peroxidases Co-Localize in Lignified Secondary Cell Walls throughout Stem Development. Plant Physiol, 184, 806–822.

Klucher, K.M., Chow, H., Reiser, L., and Fischer, R.L. (1996) The AINTEGUMENTA gene of Arabidopsis required for ovule and female gametophyte development is related to the floral homeotic gene APETALA2. Plant Cell, 8, 137–153.

Koyama, T., Furutani, M., Tasaka, M., and Ohme-Takagi, M. (2007) TCP Transcription Factors Control the Morphology of Shoot Lateral Organs via Negative Regulation of the Expression of Boundary-Specific Genes in Arabidopsis. Plant Cell, 19, 473–484.

Li, Y., Liu, C., Wang, N., Zhang, Z., Hou, L., Xin, D., et al. (2021) Fine mapping of a QTL locus (QNFSP07-1) and analysis of candidate genes for four-seeded pods in soybean. Mol Breeding, 41, 71.

Mateos, J.L., Madrigal, P., Tsuda, K., Rawat, V., Richter, R., Romera-Branchat, M., et al. (2015) Combinatorial activities of SHORT VEGETATIVE PHASE and FLOWERING LOCUS C define distinct modes of flowering regulation in Arabidopsis. Genome Biol, 16, 31.

Mizukami, Y. and Fischer, R.L. (2000) Plant organ size control: AINTEGUMENTA regulates growth and cell numbers during organogenesis. Proceedings of the National Academy of Sciences, 97, 942–947.

Monfared, M.M., Carles, C.C., Rossignol, P., Pires, H.R., and Fletcher, J.C. (2013) *The* ULT1 *and* ULT2 *trxG Genes Play Overlapping Roles in* Arabidopsis *Development and Gene Regulation*. Molecular Plant, 6, 1564–1579.

Pedersen, P. and Lauer, J.G. (2004) Response of Soybean Yield Components to Management System and Planting Date. Agronomy Journal, 96, 1372–1381.

Qadir, M., Wang, X., Shah, S.R.U., Zhou, X.-R., Shi, J., and Wang, H. (2021) Molecular Network for Regulation of Ovule Number in Plants. International Journal of Molecular Sciences, 22.

Roudier, F., Fernandez, A.G., Fujita, M., Himmelspach, R., Borner, G.H.H., Schindelman, G., et al. (2005) COBRA, an Arabidopsis Extracellular Glycosyl-Phosphatidyl Inositol-Anchored Protein, Specifically Controls Highly Anisotropic Expansion through Its Involvement in Cellulose Microfibril Orientation. Plant Cell, 17, 1749–1763.

Sangi, S., Araújo, P.M., Coelho, F.S., Gazara, R.K., Almeida-Silva, F., Venancio, T.M., and Grativol, C. (2021) Genome-Wide Analysis of the COBRA-Like Gene Family Supports Gene Expansion through Whole-Genome Duplication in Soybean (Glycine max). Plants, 10.

Schmutz, J., Cannon, S.B., Schlueter, J., Ma, J., Mitros, T., Nelson, W., et al. (2010) Genome sequence of the palaeopolyploid soybean. Nature, 463, 178–183.

Schuetz, M., Benske, A., Smith, R.A., Watanabe, Y., Tobimatsu, Y., Ralph, J., et al. (2014) Laccases Direct Lignification in the Discrete Secondary Cell Wall Domains of Protoxylem. Plant Physiol, 166, 798–807.

Scofield, S., Dewitte, W., and Murray, J.A.H. (2007) The KNOX gene SHOOT MERISTEMLESS is required for the development of reproductive meristematic tissues in Arabidopsis. The Plant Journal, 50, 767–781.

Viola, I.L. and Gonzalez, D.H. (2023) TCP Transcription Factors in Plant Reproductive Development: Juggling Multiple Roles. Biomolecules, 13.

Vogel, J.T., Liu, W., Olhoft, P., Crafts-Brandner, S.J., Pennycooke, J.C., and Christiansen, N. (2021) Soybean Yield Formation Physiology – A Foundation for Precision Breeding Based Improvement. Front. Plant Sci., 12.

Wang, Q., Zhang, W., Xu, W., Zhang, H., Liu, X., Chen, X., and Chen, H. (2024) Genome-Wide Association Study and Identification of Candidate Genes Associated with Seed Number per Pod in Soybean. International Journal of Molecular Sciences, 25.

Western, T.L. and Haughn, G.W. (1999) BELL1 and AGAMOUS genes promote ovule identity in Arabidopsis thaliana. The Plant Journal, 18, 329–336.

Yang, Z., Xin, D., Liu, C., Jiang, H., Han, X., Sun, Y., et al. (2013) Identification of QTLs for seed and pod traits in soybean and analysis for additive effects and epistatic effects of QTLs among multiple environments. Mol Genet Genomics, 288, 651–667.

Yu, Y., Li, Q.-F., Zhang, J.-P., Zhang, F., Zhou, Y.-F., Feng, Y.-Z., et al. (2017) Laccase-13 Regulates Seed Setting Rate by Affecting Hydrogen Peroxide Dynamics and Mitochondrial Integrity in Rice. Front. Plant Sci., 8.

Yuan, M., Sun, X., Yu, Z., Sun, H., Dong, S., Zhang, J., et al. (2025) QTN mapping, gene prediction, and simulation breeding of four-seed pod numbers in soybean. Front. Plant Sci., 16.

Zhong, R., Zhou, D., Chen, L., Rose, J.P., Wang, B.-C., and Ye, Z.-H. (2024) Plant Cell Wall Polysaccharide O-Acetyltransferases. Plants, 13.

Zhuang, Y., Zhao, J., Xiao, L., Liu, X., and Li, L. (2025) The LACCASE3/5/12/13 clade mediates seed coat lignin deposition and regulates imbibition and germination. Cell Reports, 44, 116584.

